# DeepPROTECTNeo: A Context-aware Personalized and Reverse Vaccinology-guided Deep Learning Framework for Immunogenicity Prediction

**DOI:** 10.1101/2025.01.04.631301

**Authors:** Debraj Das, Soumyadeep Bhaduri, Pralay Mitra

## Abstract

**Background:** The development of personalized cancer vaccines relies on accurately identifying neoepitopes capable of eliciting strong immune responses. T cell receptor (TCR)-epitope interactions are fundamental to cancer immunotherapy. Traditional computational approaches focus primarily on epitope-major histocompatibility complex (MHC) binding, often overlooking the critical contribution of TCR binding. Furthermore, the clinical applicability of existing methods is constrained by fragmented pipelines that require separate workflows for variant calling, HLA typing, and independent peptide-MHC (pMHC) or peptide-TCR (pTCR) evaluation stages.

**Results:** We present DeepPROTECTNeo, a unified deep learning framework that integrates genomic variant detection, HLA typing, high-affinity pMHC binding prediction, variant-driven TCR repertoire mining, followed by a hybrid transformer-Convolutional Neural Network dual-branch feature extractor with an explicit cross-attention-based deep learning model for TCR-epitope binding prediction. Our reverse vaccinology-inspired biologically informed architecture integrates Bidirectional Long short-term memory (Bi-LSTM) sequence features, convolutional-attention physicochemical/evolutionary descriptors via gated fusion, and TCR numbered contextual embeddings to enable residue-level interpretable modelling. Under a strict TCR-split strategy, it achieved a mean AUROC of 0.7856 and AUPRC of 0.7932 outperforming six state-of-the-art predictors by 4-5% with tight inter-fold stability. The architecture maintains high robustness against structural hard negatives and imbalanced datasets, successfully recovering 18 of 34 validated high-affinity neoepitopes from a patient-specific cancer cohort.

**Conclusions:** Experiments results demonstrate that DeepPROTECTNeo is a powerful, reliable end-to-end neoantigen prioritization framework that effectively models complex TCR-epitope interfaces directly from clinical sequencing data, providing a robust interpretable foundation to accelerate personalized cancer immunotherapy.

## 1 Background

The adaptive immune response is initiated in secondary lymphoid organs when foreign antigens, subsequent to host exposure through infectious or non-infectious means, are captured and proteolytically processed by antigen-presenting cells (APCs) into shorter peptide fragments. Upon loading onto major histocompatibility complex (MHC) molecules and presentation at the APC surface, these peptides are designated as epitopes, reflecting their role as the specific determinants that engage T cell recognition[1]. Naïve T cells interrogate peptide–MHC (pMHC) complexes through their clonotypic T cell receptors (TCRs). Productive recognition initiates intracellular signalling cascades that drive T cell activation, clonal expansion, and differentiation into effector populations[2]. The steady progress in cancer immunotherapy has leveraged these mechanisms, enabling personalized treatments where T cells target malignant cells via TCR recognition of tumour-derived peptides presented on MHC molecules[3–5]. In particular, the identification of tumour-specific neoantigens, i.e., peptides arising from somatic mutations exclusive to cancer cells and absent from normal tissues, has transformed personalized therapeutic strategies[6,7]. Unlike tumour-associated antigens (TAAs), neoantigens are uniquely expressed by cancer cells, making them highly attractive targets for vaccine development and adoptive T cell therapies[7–9].

Next-generation sequencing (NGS) techniques, particularly whole-exome sequencing (WES) and whole-genome sequencing (WGS), enable us to capture tumour-specific somatic variants followed by RNA Sequencing to confirm their expression[10,11]. Translating these raw data into clinically actionable targets, however, requires a multi-step workflow including precise HLA genotyping (e.g., ArcasHLA[12], OptiType[13]), in silico pMHC binding predictions (e.g., NetMHCpan[14], MHCflurry[15]), followed by T-cell receptor (TCR) peptide (pTCR) binding, which is often neglected by traditional MHC-peptide binding approaches[16,17]. Despite these advances, the clinical applicability of existing methods is mostly confined to multiple input files at different stages of processing and separate workflows for TCR profiling, HLA typing, and variant calling[16,18], pMHC and pTCR binding, thereby constraining clinical applicability.

Recently, various deep-learning frameworks have been developed for TCR-epitope binding prediction. Still, substantial challenges remain in aligning these computational advances with the nuanced realities of immunobiology and clinical implementation. Early CNN models like ImRex[19], NetTCR-2[20], and TITAN[21] primarily focus on local motifs, often failing to capture long-range dependencies. Recurrent and autoencoder-based frameworks, including ERGO-AE[22], DeepTCR[23], focus on mapping CDR3β and peptide sequences into meaningful latent spaces, yet they offer only marginal gains in zero-shot settings. The introduction of transformer-based approaches like ATM-TCR[24] and TEPCAM[25] yields strong in-domain accuracy but still reverts to near-random predictions on truly novel antigens. Hybrid pMHC-TCR models such as pMTnet[26] and TCRGP[27] added peptide-MHC binding priors and even docking-derived constraints, limited by the dependency on paired α/β-chain and HLA information. Additionally, a wave of interpretable and hybrid-learning frameworks for neoantigen immunogenicity prediction has combined attention-based mechanistic explanations with multi-modal physicochemical, structural, and sequence feature fusion to improve both predictive accuracy and biological transparency[28–30]; however, most of these frameworks remain restricted to either MHC-stage neoantigen filtering or stand-alone TCR–epitope binding scoring, without unifying both stages within a single clinically deployable pipeline. A parallel progress in self-supervised pre-training produced TCR-specific language models like BertTCR[31] and STAPLER[32] trained on tens of millions of unlabelled sequences, alongside LSTM-based CatELMo[33], yet they face trade-offs between generality and specificity. Most recent deep-learning-based immunogenicity prediction efforts have further expanded this methodological space through transformer and graph-based residue representations, contextualised pretrained encoders, and multi-task supervision for TCR– neoepitope binding[34,35], collectively reinforcing that joint sequence, structure, and immunogenicity modelling is an actively evolving direction in computational immunology, yet integration of these advances into an end-to-end neoantigen prioritisation pipeline remains an open problem. Given this development in DL-based techniques, the ultimate challenge in TCR-peptide specificity modelling arises from the CDR3β loop itself: it mediates the majority of peptide contacts yet exhibits staggering sequence diversity, context-dependent binding modes, and, in most high-throughput assays, lacks paired α-chain information[36,37].

While these prior TCR-epitope binding frameworks individually advance either sequence-based prediction (ATM-TCR[24], TEPCAM[25], NetTCR[20]), pMHC-aware modelling (pMTnet, TCRGP), or large-scale self-supervised pre-training (BertTCR[31], STAPLER[32]), none jointly integrate (i) residue-level rotary cross-attention between CDR3β and epitope, (ii) a physicochemical and evolutionary descriptor branch fused through a learned gate with the sequence representation. Our framework is conceptualised around this tri-partite novelty. DeepPROTECTNeo is a unified pipeline for personalized neoantigen prioritization streamlining the entire workflow from processing raw RNA-Seq reads through somatic variant calling, HLA inference, peptide–MHC filtering, and downstream TCR-epitope binding analysis (Fig. 1). At its core, DeepPROTECTNeo employs a dual-branch, context-aware Transformer-CNN architecture: a sequence branch leveraging bidirectional LSTMs with learned positional encodings to capture long-range dependencies in peptide and TCR CDR3β sequences, and a convolutional branch encoding 102 physicochemical and BLOSUM-derived descriptors via deep residual networks with channel-wise self-attention (Fig. 2A, B). A gated fusion module integrates these complementary representations, while a cross-attention mechanism with rotary embeddings captures fine-grained residue-level interactions, enabling both robust prediction of TCR-epitope binding and interpretability by recovering key anchor residues and novel contact motifs and progressively capturing a clear separation through propagation across different layers (Fig. 2C). Beyond achieving state-of-the-art predictive performance across multiple benchmarking cohorts, DeepPROTECTNeo demonstrates clinical relevance when evaluated on the TESLA consortium’s experimentally validated neoepitope dataset, where it consistently separates binders from non-binders and recovers the majority of confirmed epitopes. These results establish DeepPROTECTNeo as a powerful framework to accelerate translational pipelines for patient-specific vaccine and adoptive T-cell therapy design, as detailed in the sections that follow.

**Figure 1.**
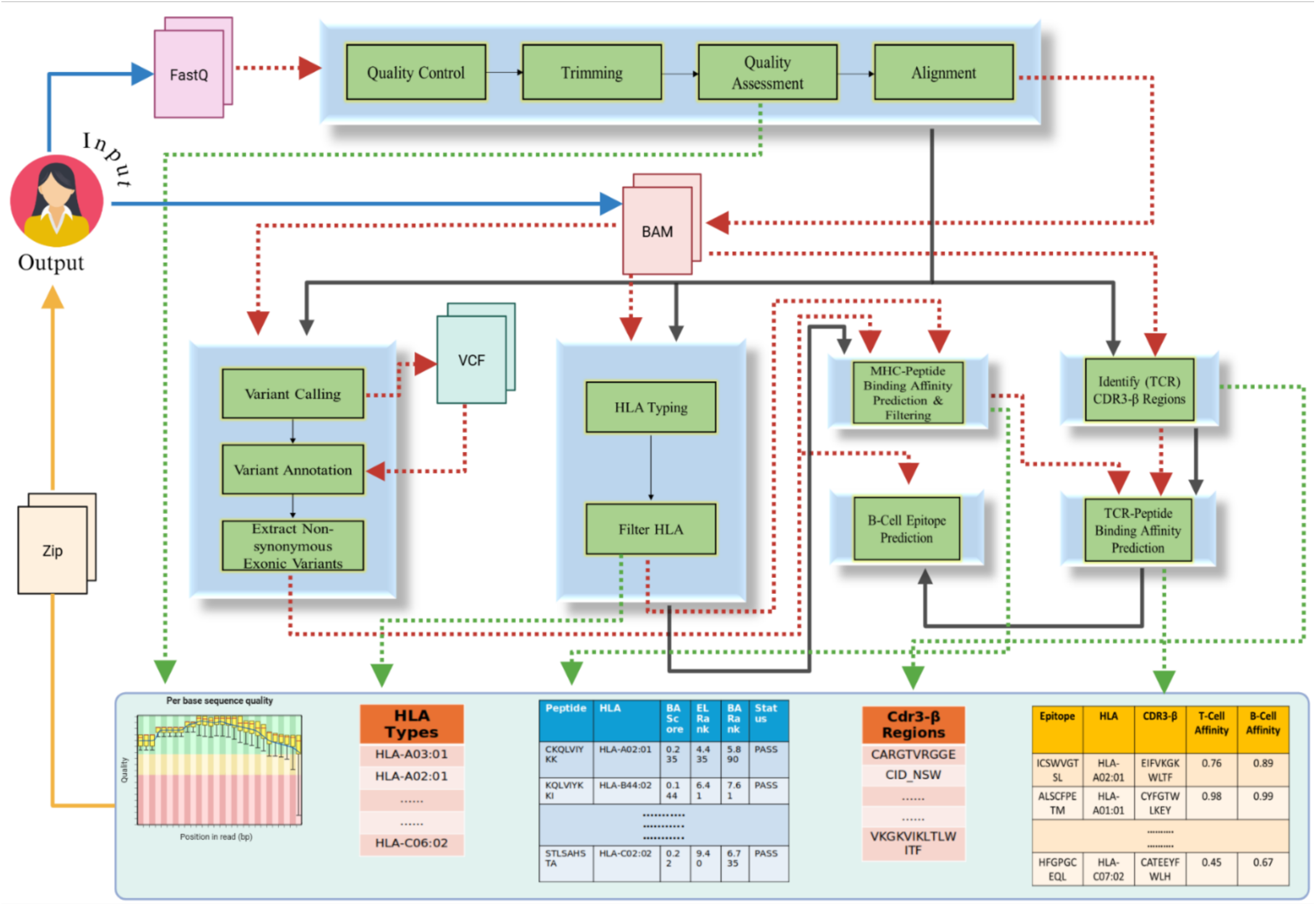
Integrated DeepPROTECTNeo neoantigen discovery workflow. Solid blue arrows denote user inputs (raw FASTQ or BAM or candidate CDR3β–epitope pairs); solid black arrows show core processing steps—QC/trimming/alignment → variant calling & annotation → HLA typing → CDR3β extraction. Candidate peptides are scored for MHC binding (NetMHCpan) and B-cell epitopes, then matched with CDR3β sequences for DeepPROTECTNeo binding prediction. Red dashed lines trace intermediate data transfers (VCFs, HLA calls, CDR3β lists); green dashed lines indicate final data outputs (QC reports, HLA alleles, neoepitope and repertoire tables, binding scores); solid orange arrow marks the ZIP archive delivered to the user.

**Figure 2.**
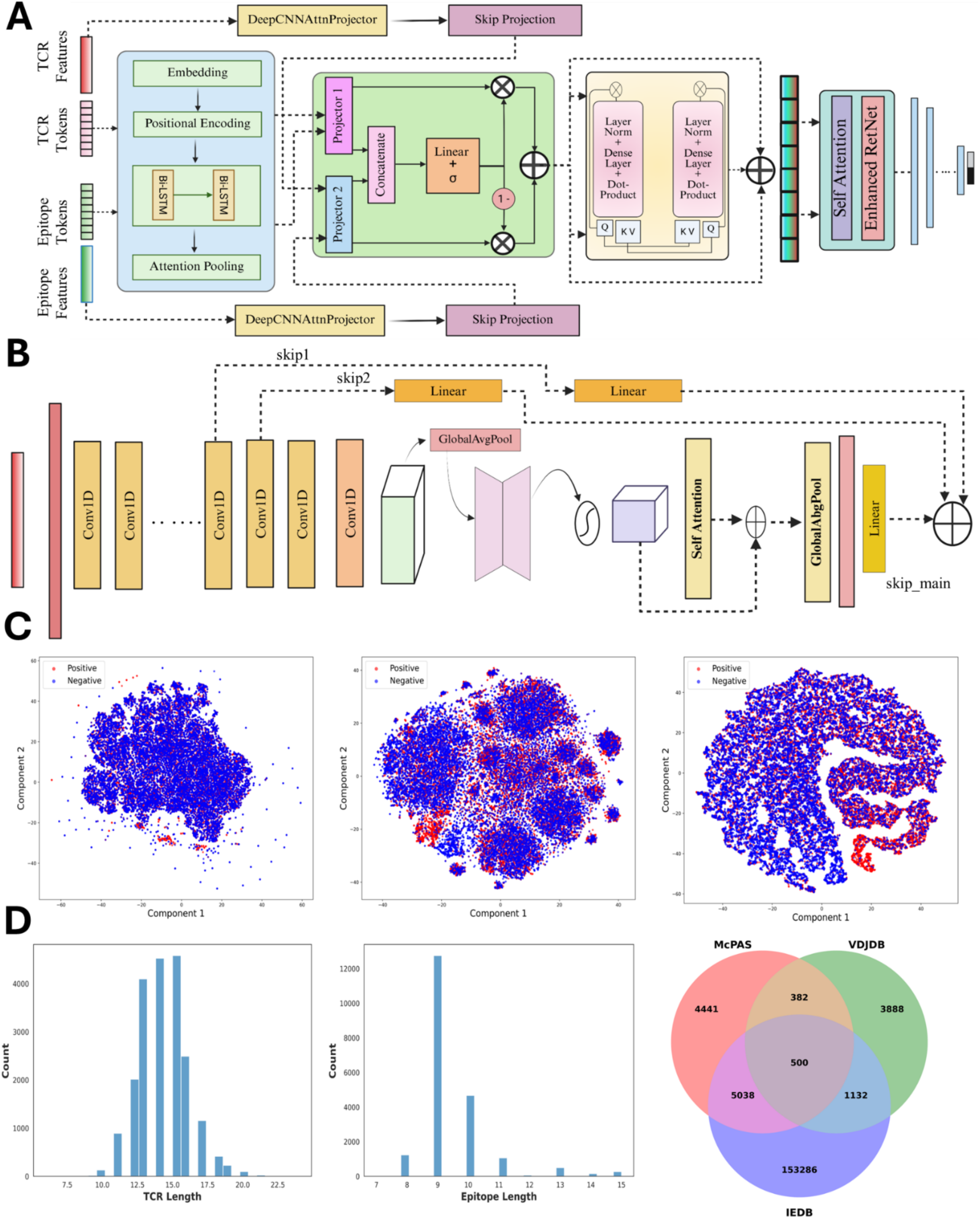
DeepPROTECTNeo architecture, latent-space embeddings, and dataset. **(A**) Paired TCR (20 aa) and epitope (11 aa) tokens plus 102-dim physicochemical descriptors enter two streams: a token branch that embeds into ℝ^(B×L×240), applies learned positional encodings, a 2-layer BiLSTM and attention pooling to yield 240-dim features; and a skip branch that runs 32 Conv1D + Channel Attention (CA) blocks (C=64, ks=5) with optional self-attention over descriptors, pools to a 60-dim vector, extracts two intermediate maps and t_final into three 30-dim skip vectors, and gates them into a single 30-dim summary. A GatedFusion per side produces t_fused_, p_fused_ ∈ ℝ^(B×240)^; a 6-head rotary cross-attention yields x_fused_ ∈ ℝ^(B×240)^; concatenation [t_fused_; p_fused_; x_fused_] ∈ ℝ^(B×720) is refined by 6-head self-attention plus a 24-layer ReZero RetNet and classified by a 2-way MLP. (**B)** Skip branch detail. Shows Conv1D block outputs (B×64×51), 1×1 Conv→CA, global average pooling to t_final_ (B×60), two intermediate skip-map projections and t_final projections to 30-dim, concatenation to 90 dim, and gating to 30 dim. (**C)** t-SNE embeddings. Positive (red) vs negative (blue) test pairs projected into 2D using features from (left) concatenated fusion vector [t_fused_; p_fused_; x_fused_] before extra self-attention, (middle) cross-attention x_fused_ before extra self-attention, and (right) final post-RetNet outputs demonstrating progressively clearer class separation. (**D)** Histogram of TCR β-chain CDR3 lengths distribution (left), histogram of epitope sequence lengths (middle), and Venn diagram of unique TCR–epitope pair overlap among IEDB (Class I & II), McPAS, and VDJdb (right).

## 2 Results

### 2.1 DeepPROTECTNeo Improves Binding Performance on Strict TCR Split Data

We investigated DeepPROTECTNeo’s CDR3β-epitope binding prediction capabilities using a stringent TCR-split strategy that mirrors authentic clinical immunotherapy scenarios. We curated a unified dataset (Additional File 1, Note S1; Additional File 2, Figure S1) consisting of 140992 unique CDR3β-epitope interacting pairs among 130303 diverse TCR repertoire and 1903 epitopes, a source-wise data distribution is given in Additional File 3, Table S1 (see CDR3β and epitope length distribution and their overlap across the datasets in Fig. 2D). Finding experimentally validated non-binding interaction pairs possesses a significant challenge in this domain, we relied on computational strategy to overcome this by generating negative pairs through an epitope shuffling process while preserving underlying sequence distribution characteristics maintaining biological plausibility without introducing systematic biases. The paired TCR-epitope data were split into 5 equal, mutually disjoint folds based on TCR sequences. For benchmarking, we carefully selected six state-of-the-art, single-chain TCR-epitope predictors, including ATM-TCR[24], ERGO-AE[22], ERGO-LSTM[22], TEINet[38], epiTCR[39] and NetTCR-2.0[20], based on their ability to operate using only CDR3β input, thus matching DeepPROTECTNeo’s single-chain paradigm (See Additional File 1, Note S2; Additional File 2, Figure S2 for details about training and performance across epochs). To ensure a fair and rigorous comparison, we retrained each of these models from scratch on our unified dataset using their default parameter settings.

We evaluated all models using key classification metrics Balanced Accuracy (BA), macro-averaged F1-score, AUROC, and AUPRC (See Additional File 1, Note S3 for evaluation metric’s details) to capture both class-wise prediction quality and overall ranking performance. In particular, BA was preferred over raw accuracy because of its ability to equally weight true positive and true negative rates, crucial for the wide distribution of TCR sequences in the training data. Combined ROC and precision–recall analyses (Fig. 3A, B) reveal that DeepPROTECTNeo achieves a mean±s.d. AUROC of 0.7856±0.0023 and AUPRC of 0.7932±0.0024, significantly outperforming all other models by a margin of 4 percentage point gain in both metrics. Strong early-retrieval performance, evidenced by high precision at the leftmost (low-recall) region of the precision-recall curves, suggests that the top-ranked predictions are densely populated with true binders. Moreover, the narrow-shaded bands (mean±1s.d. across five TCR-split folds) quantify consistent performance over the folds: DeepPROTECTNeo’s inter-fold s.d. is just 0.0023 for AUROC and 0.0024 for AUPRC, whereas other models, particularly epiTCR, show high variability (0.1203 for AUROC and 0.1240 for AUPRC), suggesting inter-fold performance fluctuations, making its top predictions highly unreliable. Paired two-tailed t-tests confirm that DeepPROTECTNeo’s improvements for both the metrics over ATM-TCR, ERGO-AE, ERGO-LSTM, TEINet, and NetTCR-2 are highly significant (p<1×10⁻⁴).

**Figure 3.**
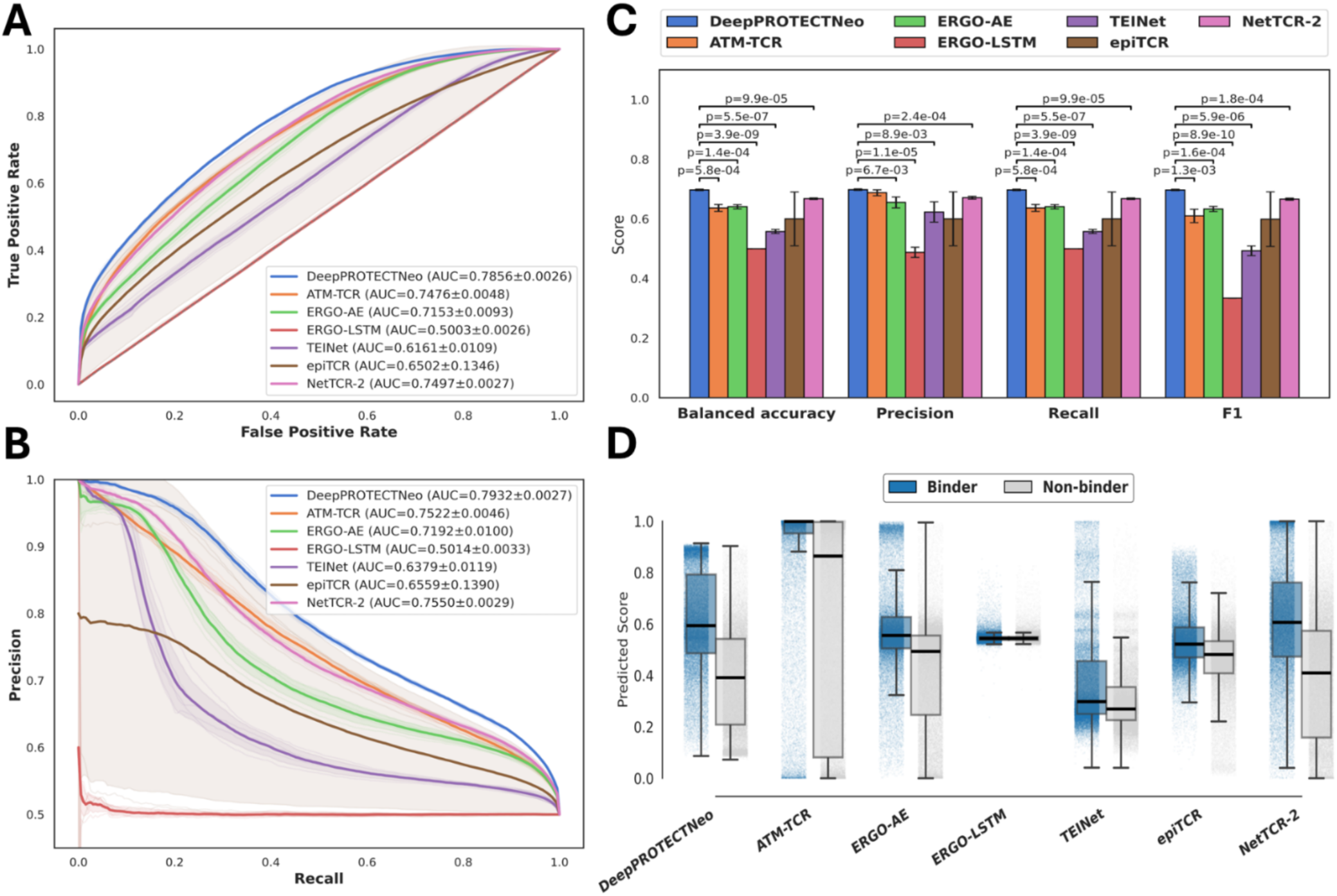
Benchmarking DeepPROTECTNeo against CDR3β-only TCR–epitope predictors under stringent TCR-split over 5 folds. **A)** Mean receiver-operating-characteristic (ROC) curves (true-positive rate versus false-positive rate) for DeepPROTECTNeo (blue), ATM-TCR (orange), ERGO-AE (green), ERGO-LSTM (red), TEINet (purple), epiTCR (brown), and NetTCR-2 (pink) computed over five independent folds. Shaded bands denote ±1 s.d. across folds; legend reports mean AUROC ± s.d. **B)** Mean precision–recall (PR) curves (precision versus recall) for the same models and folds, with shaded bands = ±1 s.d. and mean AUPRC ± s.d. in the legend. **C)** Fold-averaged balanced accuracy, macro-precision, macro-recall, and macro-F1 at a fixed decision threshold. Bars = mean over five folds; error bars = ±1 s.d. Pairwise significance versus DeepPROTECTNeo was determined by paired two-tailed t-tests (exact p-values shown for significant ones). **D)** Distribution of per-interaction predicted scores for binder (blue) and non-binder (grey) classes across all models. Boxplots show median (central line), 25th–75th percentiles (box limits) and 1.5× IQR whiskers; overlaid jittered points (30% subsample) illustrate score density. All analyses are based on a unified dataset of 140,992 unique CDR3β–epitope pairs evaluated under five TCR-split folds.

To complement the continuous-ranking evaluation in earlier, we demonstrated a threshold-based assessment by showing fold-averaged metrics under a uniform classification cutoff (Fig. 3C). DeepPROTECTNeo achieves a BA of 0.6973±0.0020, outperforming the next best (NetTCR-2: 0.6678±0.0025) by 4.4 %; a precision of 0.6981±0.0021, a 0.8 % gain over closest ATM-TCR (0.6923±0.0100); a recall of 0.6973±0.0020, around 3% above NetTCR-2 (0.6678±0.0025); and an F1 of 0.6969±0.0020, 3.1 % higher than NetTCR-2 (0.6662±0.0030).

All pairwise tests for BA, recall, and F1 show highly significant improvements over ATM-TCR (p=5.8×10^-4^ to 1.3×10^-3^), ERGO-AE (p=1.4×10^-4^ to 1.6×10^-4^), ERGO-LSTM (p=8.9×10^-10^ to 3.9×10^-9^), TEINet (p=5.5×10^-7^ to 5.9×10^-6^), and NetTCR-2 (p=9.9×10^-5^ to 1.8×10^-4^). Precision gains were likewise significant versus ERGO-AE (p=6.7×10^-3^), ERGO-LSTM (1.1×10^-5^), TEINet (8.9×10^-3^), and NetTCR-2 (2.4×10^-4^). We demonstrated the raw score distributions for binders against non-binder pairs (Fig. 3D), and the Coefficient of Variation (CV) affirms that DeepPROTECTNeo yields a clear gap between binder (mean 0.611±0.193, CV 0.316) and non-binder (mean 0.385±0.194, CV 0.505) predictions with moderate inter-fold variability. Whereas, ATM-TCR, despite having a high binder (mean 0.885±0.257, CV 0.290), lacks clear non-binder scores (mean 0.614±0.422, CV 0.688), while ERGO-LSTM shows zero discrimination (both classes mean≈0.547±0.02, CV≈0.036) and TEINet’s binder scores fluctuate widely (CV 0.512 for binder and 0.4 for non-binder). NetTCR-2 also displays high variability in both classes (CVs 0.373/0.637). Although epiTCR achieves the lowest CVs (0.215/0.285), its small mean separation (0.539 vs 0.454) hinders its discriminative power. Together, these results demonstrate that DeepPROTECTNeo achieves the best balance of strong binder, non-binder separation, and consistent score stability across folds.

To assess the robustness of the decision boundary against realistic clinical non-binding distributions, we constructed two additional evaluation datasets. First, a structural Hard-Negatives set was built by clustering all unique training CDR3β sequences using CD-HIT at an 85% sequence-identity threshold; 108,543 non-binding pairs were then synthesised by pairing each epitope with non-cognate TCRs drawn from the same identity cluster as a confirmed biological binder, thereby enforcing strict structural homology between negatives and true binders. Second, a 1:2 positive-to-negative imbalanced set (338,382 interactions) was constructed to reflect realistic clinical prevalence. On the Hard-Negatives set, DeepPROTECTNeo achieved AUROC = 0.7546 ± 0.0019, with a ΔAUROC of 0.0310 (3.95%) below the 0.7856 ± 0.0023 baseline. On the 1:2 imbalanced set, AUROC was preserved at 0.7883 ± 0.0028 with a normalised AUPRC lift of +0.346 above random (vs. +0.293 at 1:1). Together, these results confirm that DeepPROTECTNeo has learned underlying biochemical specificity rules rather than compositional biases of the random-recombination negatives, and that the decision boundary generalises to both high-similarity distractors and deeply imbalanced clinical screening conditions.

### 2.2 Contributions of Architectural Components on Binding Predictions

We have shown a detailed module-wise model parameters, their input-output dimension along with a complete block by block by architecture in Additional File 2, Figure S3 and module-wise parameter details in Additional File 3, Table S2. To find the best model configuration, we performed a grid search for achieving a stable model configuration over a range of hyperparameters (see Additional File 3, Table S3), followed by sensitivity analysis revealing its robustness to hyperparameter variations, with only modest fluctuations across top configurations. All primary metrics (F1, accuracy, precision, recall) showed minimal variance (standard deviation≤0.004), highlighting architectural stability. The skip connection dimension (skip_dim) exhibited the strongest positive correlation with all performance metrics (Spearman ρ≈0.40-0.45), confirming its importance for effective information transfer and gradient flow (see Fig. 4). CNN embedding dimension (cnn_dim) and the number of CNN blocks had moderate positive effects, supporting the extraction of multi-scale physicochemical features. The dimensionality of the stacked CNN 1D residual blocks, and the CNN dimension shows its superiority over the other model parameters when compared on validation loss, showcasing its importance in projecting the features to the forward layers, which is further confirmed using F scores derived using ANOVA (Additional File 2, Figure S4). The removal of the feature-enriched skip connections led to a drop in accuracy and F1, confirming their role in stabilizing learning and preventing feature degradation, albeit with a slight trade-off in AUROC and AUPRC. Removal of cross-attention resulted in a minor decrease in all metrics, underlining its value for modelling inter-sequence dependencies. The complete model, integrating both skip connections and cross-attention, consistently achieved the highest scores across all evaluation metrics (ablation analysis in provided in Additional File 3, Table S4). DeepPROTECTNeo architecture demonstrates strong resilience to hyperparameter changes, with skip connections and cross-attention emerging as the most influential components for maximizing predictive accuracy and generalization. The model’s design ensures reliable and stable performance across a range of settings, validating its suitability for robust TCR-epitope binding prediction in diverse clinical and biological contexts.

**Figure 4.**
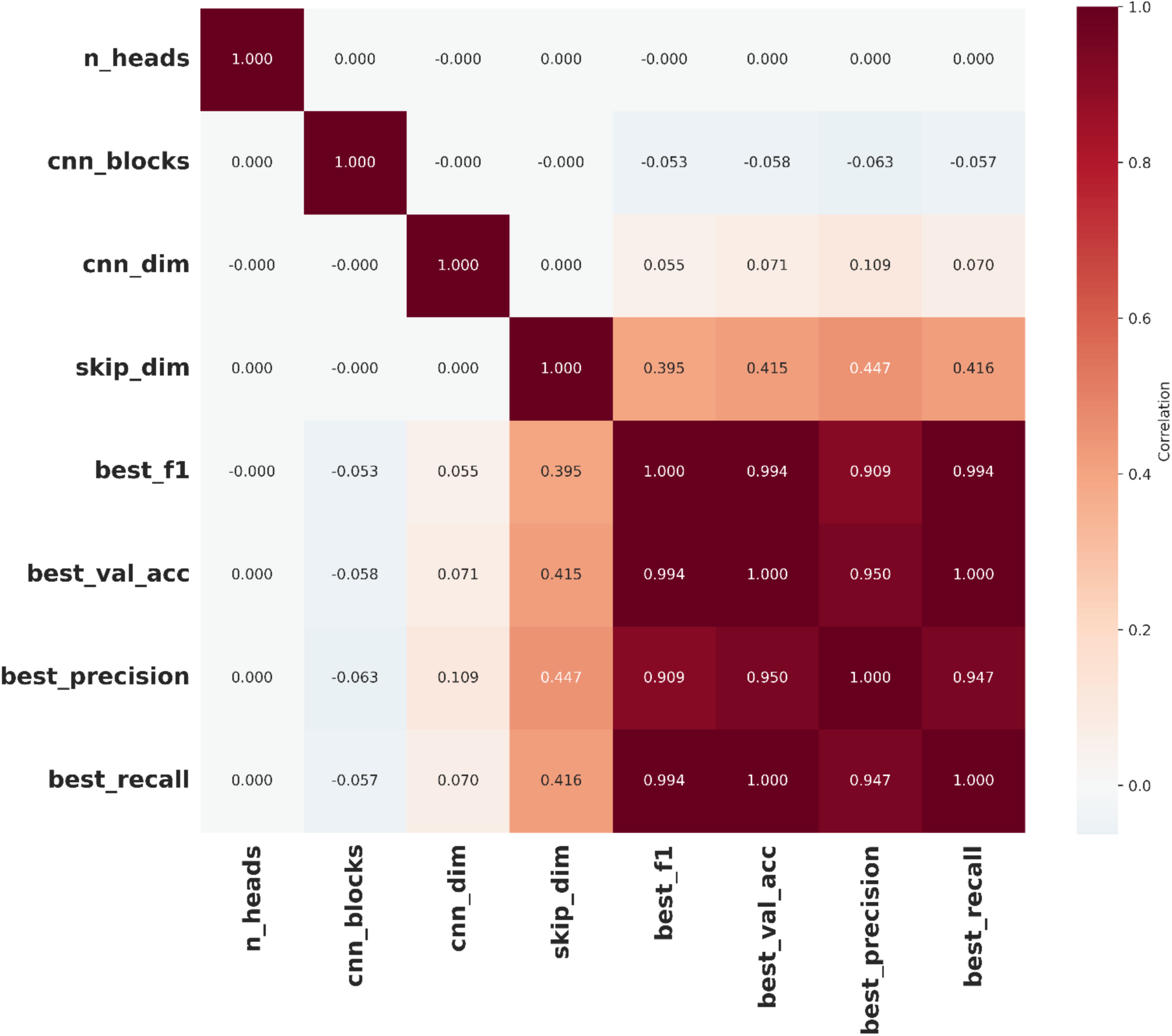
Correlation heatmap displaying relationships between model hyperparameters (n_heads, cnn_blocks, cnn_dim, skip_dim) and performance metrics (best_f1, best_val_acc, best_precision, best_recall), highlighting strong positive correlations between key evaluation scores and skip_dim, while revealing little or no correlation between hyperparameters and performance outcomes. The colour scale represents the degree of linear association, with darker shades indicating higher correlations.

### 2.3 DeepPROTECTNeo Demonstrates Improved Binding Performance on SARS-CoV-2 data

To evaluate DeepPROTECTNeo’s generalizability to real-world TCR-epitope diversity, we compared its performance on the ImmuneCODE[40] dataset against the six binding models as discussed earlier. ImmuneCODE is a repertoire of experimentally validated TCR-epitope interaction pairs derived from over 1000 SARS-CoV-2-infected individuals. All six models were trained on our unified dataset over 5 stratified folds to retain the uniformity of the predictions. To prevent data leakage, we removed all the interacting pairs from ImmuneCODE that were also present on our unified dataset, finally retaining 18811 interactions. Negative pairs (non-binding TCR-epitope pairs) were generated using random recombination of the positive pairs, yielding a balanced 1:1 dataset for performance evaluation. DeepPROTECTNeo consistently outperforms state-of-the-art baselines across all key metrics (Fig. 5A). Aggregated over five folds, we achieved the highest BA (0.6237±0.0042), F1-macro (0.6223±0.0039), AUROC (0.6752±0.0047) and AUPRC (0.6766±0.0049), with paired *t*-test confirming significant gain over nearest best performing model for all the metrices: BA (ATM-TCR; Score: 0.5664±0.0179, p=4.57×10^-3^), F1-macro (ERGO-AE: 0.5258±0.0298, p=3.92×10^-3^), AUROC (ATM-TCR; Score: 0.6623±0.0077, p=8.74×10^-3^), AUPRC (ATM-TCR; Score: 0.6590±0.0049, p=1.93×10^-3^). Furthermore, DeepPROTECTNeo exhibits the lowest cross-fold variability (e.g., AUROC: σ=0.0047 vs the closest ATM-TCR σ=0.0077), demonstrating its robust generalization over the fold in a zero-shot setting. Due to masking mis-calibration of high-aggregated scores[41], we further evaluated the reliability of our binding predictions (Fig. 5B).

**Figure 5.**
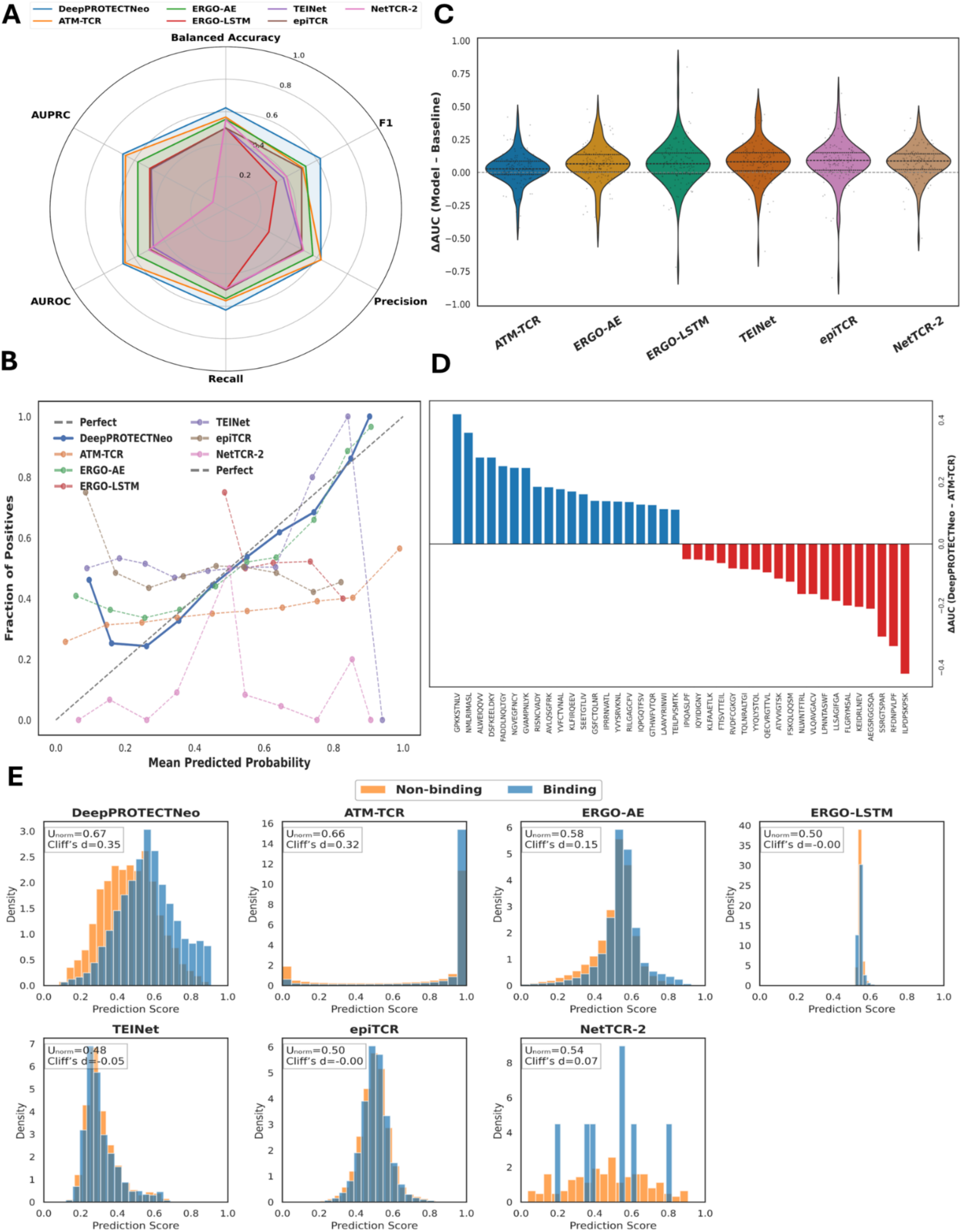
Performance evaluation of DeepPROTECTNeo on the ImmuneCODE benchmark. **A)** Radar plot of six key metrics: balanced accuracy, F1, precision, recall, AUROC and AUPRC, showing the fold-averaged mean for DeepPROTECTNeo (blue) and six comparator models. **B)** Reliability (calibration) diagram: observed binder fraction versus mean predicted probability in ten uniform bins for each model; the gray dashed line is perfect calibration. **C)** Violin plots of per-epitope ΔAUC (DeepPROTECTNeo minus each baseline) with individual points and median (dashed line). **D)** Waterfall of the 20 epitopes with the largest ΔAUC gains (blue; median +0.174, IQR 0.111, max +0.420) and losses (red; median −0.143, IQR 0.120, min −0.420) compared to the closest ATM-TCR. **E)** Overlaid histograms of prediction scores for true binders (blue) and non-binders (orange), annotated with normalized Mann–Whitney U (Uₙₒᵣₘ = U / (m·n), where U is the test statistic and m, n are sample sizes) and Cliff’s δ, illustrating DeepPROTECTNeo’s superior class separation (Uₙₒᵣₘ = 0.67, d = 0.35).

DeepPROTECTNeo’s curve closely follows the diagonal line more tightly than any competitor, yielding the lowest Brier score of 0.2267 and minimal Expected Calibration Error (ECE) of 0.0218. Using the Hosmer–Lemeshow test, DeepPROTECTNeo’s χ² statistic (836) was recorded orders of magnitude smaller than all the baselines (χ² score>1700 for every other model), demonstrating its superior probability estimation. We examined per-epitope ΔAUC across all epitopes with violin plots (Fig. 5C), capturing the full range of gains and losses over all the epitopes. For every model, the one-sample t-test rejects the null hypothesis of zero mean difference against ATM-TCR (mean ΔAUC=+0.027, IQR=0.101; p=1.03×10⁻²), ERGO-AE (+0.065, IQR=0.131; p=1.09×10⁻⁶), ERGO-LSTM (+0.066, IQR=0.158; p=1.61×10⁻⁵), TEINet (+0.080, IQR=0.138; p=2.13×10⁻⁸), epiTCR (+0.091, IQR =0.133; p=1.48×10⁻⁷) and NetTCR-2 (+0.086, IQR=0.119; p=2.85×10⁻¹⁰). Building on the bi-directional consistency in Fig. 5C, 40-epitope waterfall (Fig. 5D) reveals that DeepPROTECTNeo markedly outperforms its closest competitor, ATM-TCR, on its top 20 gains (median ΔAUC=+0.174, IQR=0.111; maximum +0.420), while its 20 worst peptides show a median loss of only –0.143 (IQR=0.120; minimum –0.420, a near-symmetric profile that highlights widespread, consistent improvements. Fig. 5E shows DeepPROTECTNeo’s raw score distributions, highlighting its smooth, well-calibrated confidence and superior binder/non-binder separation for reliable thresholding. Our binding prediction model demonstrated the largest separation between true binders and non-binders (Cliff’s δ=0.35, U test, p≈0.00), with scores distributed almost symmetrically around the decision boundary (Skew test; skewness score=0.071, p=5.64×10^-36^), indicating only a minimal right-tail bias. In contrast, ATM-TCR achieved a similar effect size (δ=0.32) but exhibited pronounced left skew (skewness score=–1.731, p<10^-36^), with nearly half of the predictions saturating at the upper limit, limiting threshold resolution. ERGO-AE showed moderate discrimination (δ=0.15) and mild left skew (skewness score=–0.852, p≈0.00), while ERGO-LSTM, epiTCR, and TEINet yielded effect sizes near zero, indicating negligible separability. NetTCR-2 displayed only a marginal effect (δ=0.07) and uneven binder score distribution (skewness score=–0.049, p=0.798). These comparisons highlight that our framework not only quantifies relative ranking ability but also characterises the underlying score distribution, revealing model-specific biases that influence threshold-based performance.

### 2.4 DeepPROTECTNeo Outperforms Baselines on Mutation and Viral Binding Data

Next, we investigated the performance of our model on two challenging TCR-epitope interaction datasets, namely Mutation and Viral. Mutation data is sourced from studies by the ePytope-TCR[42,43] team, and it comprises 4244 TCR-epitope pairs containing six TCRs against 132 single amino acid mutations of the neo-epitope *VPSVWRSSL*, and 172 mutations of the human CMV epitope *NLVPMVATV* in which experimentally validated epitopes were systematically altered by single-amino-acid substitutions at key positions. Notably, this dataset comprises three TCR repertoires with annotated activation scores through exchanging the amino acid residues and providing a continuous activation score using NFAT (Nuclear Factor of Activated T cells) reporter expression in flow cytometry for each combination of TCR-epitopes. The viral dataset consists of 8932 TCR 9-mer epitope interactions, sourced from the same study by ePytope-TCR[43], originated from a vaccine cohort study by Kocher et al.,[44] and the BEAM-T pipeline example datasets provided by 10x Genomics, originally drawn from SARS-CoV-2, HHV-1, EBV, and IAV from 14 donors. For benchmarking, the same five-fold strategy was followed here for all the models. In the Mutation dataset, DeepPROTECTNeo emerges as the top-performing model (Fig. 6A) on every key metric, achieving the highest BA (0.5634±0.0183), AUROC (0.5978±0.0153), AUPRC (0.3562±0.0102), and F1-macro (0.5626±0.0172). The closest competitor in balanced accuracy is ATM-TCR (0.5368±0.0163), while ERGO-AE registers the second-highest AUROC (0.5678±0.0258) and F1-macro (0.5328±0.0191), and ATM-TCR is the closest in terms of AUPRC (0.3338±0.0203). For the Viral dataset (Fig. 6B), DeepPROTECTNeo leads in BA (0.5656±0.0191) and AUROC (0.6091±0.0202), with ATM-TCR being a close second in BA (0.5602±0.0060) and NetTCR-2 in AUROC (0.6029±0.0262). We also compared our model’s performance based on the given trained weights for different categories of models (including pMHC-TCR, epitope-TCR α/β models) following the strategy and metrics provided by ePytope-TCR[43] (Additional File 3 Table S5, S6)[19,26,32,45–52], and observed that DeepPROTECTNeo emerged as one of the top-performing models on both of these datasets in every metric.

**Figure 6.**
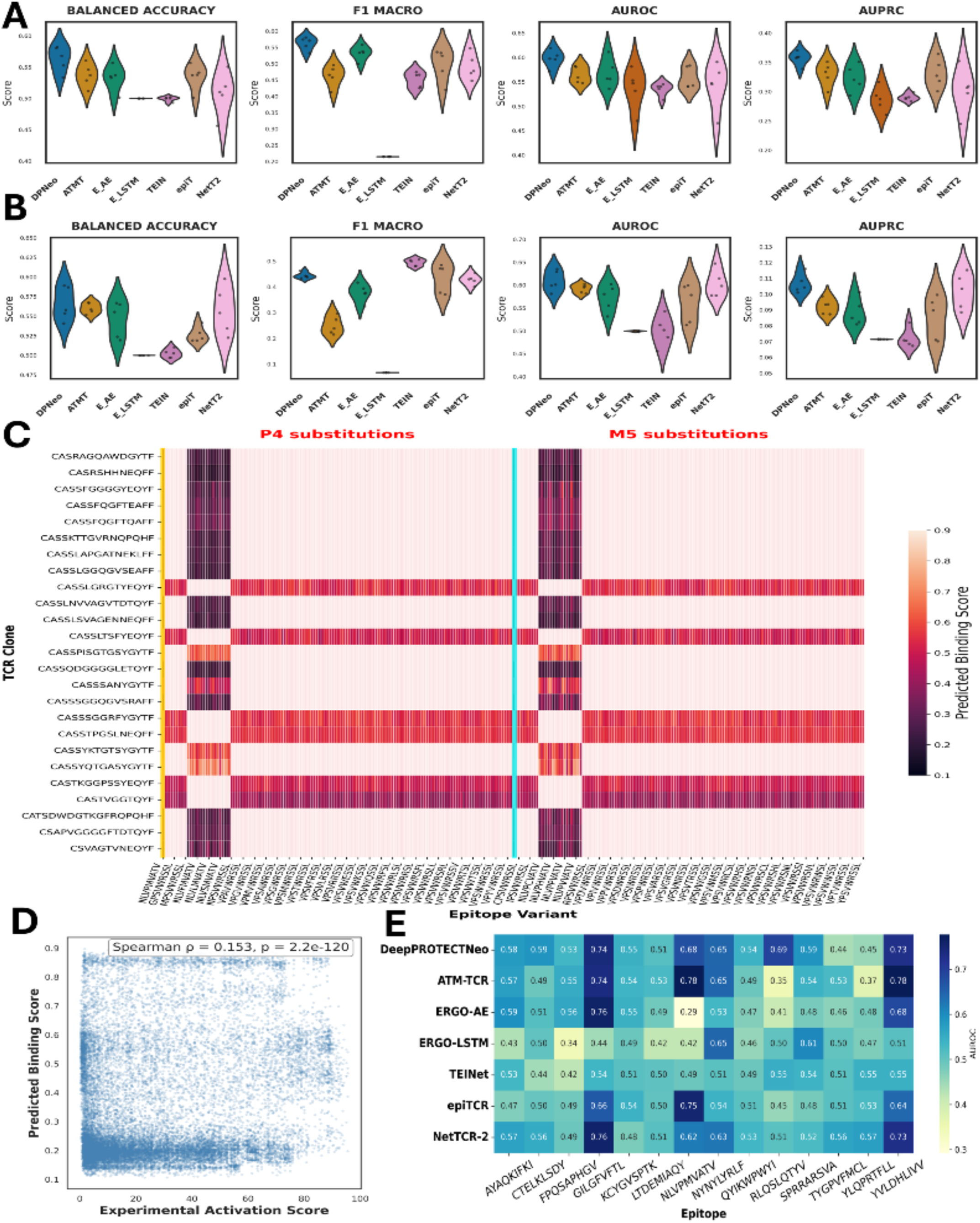
Comprehensive Performance Assessment and Biophysical Insights of DeepPROTECTNeo on Mutation and Viral datasets. **A)** Violin plots of five-fold balanced accuracy, F1-macro, AUROC and AUPRC on the engineered mutation dataset (4244 TCR– peptide pairs across 26 TCR clones). DeepPROTECTNeo (DPNeo) is shown as the leftmost (blue) violin in each panel, followed by ATM-TCR (ATMT), ERGO-AE (E_AE), ERGO-LSTM (E_LSTM), TEINet (TEIN), epiTCR (epiT) and NetTCR-2 (NetT2); individual fold scores are overlaid as black dots. **B)** Corresponding performance on the Viral dataset (8932 TCR–epitope interactions from CMV, EBV, and influenza). **C)** Two-anchor mutation-scan heatmap on Mutation data: DeepPROTECTNeo’s mean predicted binding for every P4 (left) and M5 (right) substitution across 26 TCR clones. Wild-type columns are outlined in gold (P4→P) and cyan (M5→M); colour scale from low (dark) to high (pale) affinity. **D)** Scatterplot of predicted binding versus experimental activation scores on Mutation data, with Spearman’s ρ = 0.153 and two-tailed p = 2.2 × 10⁻¹²⁰ indicated, demonstrating a significant monotonic relationship. **E)** Epitope-wise AUROC heatmap on Viral data: five-fold average AUROCs for seven models across fourteen viral 9-mer peptides; deeper blue denotes superior discrimination.

To dissect the biophysical pattern encoded by DeepPROTECTNeo, we performed an exhaustive single-residue scan at the two central “hotspot” positions P4 and M5 across 20 amino-acid variants for each of 26 experimentally characterized TCRs on Mutation data (Fig. 6C). Each block is anchored by the unmodified peptide (gold border at P4→P; cyan border at M5→M) to serve as an internal baseline. In the P4 block, a subset of “P4-tolerant” clones (e.g., *CASSYQTGASYGYTF*, *CASSTPGSLNEQFF*) retains uniformly high scores (≈0.7-0.9) even when Proline is replaced by charged or bulky residues, whereas “P4-sensitive” receptors (e.g., *CASSFGQGVTQYF*, *CASTVGGTLQYF*) collapse to <0.2 upon any deviation from Proline, revealing strict dependence on that hydrogen-bonding donor. Conversely, in the M5 block charged or polar swaps (Asp, Lys) uniformly abrogate binding (scores<0.2), while hydrophobic substitutions (Val, Leu, Ile) preserve or enhance affinity (≈ 0.6–0.8), recapitulating the classical hydrophobic TCR pocket. Strikingly, many clones that tolerate P4 substitutions become exquisitely specific at M5 (and vice versa), showing that even adjacent side chains sculpt distinct complementarity-determining surfaces. We computed the Spearman rank correlation between model scores and experimentally measured activation scores (binarized by the thresholds of 66.09% and 40.0% for the neo-antigen and CMV datasets, as given in the source study[43]) across all TCR-epitope pairs (Fig. 6D) on the Mutation data.

DeepPROTECTNeo achieved the strongest monotonic relationship, Spearman’s ρ=0.157 (p=2.4×10^-125^), outperforming every other competing method: ERGO-AE (ρ=0.116), ATM-TCR (ρ=0.098), NetTCR-2 (ρ=0.102), epiTCR (ρ=0.074), TEINet (ρ=0.071), and ERGO-LSTM (ρ=0.072). Further, we evaluated how DeepPROTECTNeo generalizes across diverse viral targets by computing per-epitope AUROCs for fourteen distinct 9-mer peptides on Viral data. In the heatmap (Fig. 6E), each row represents a model, while each column represents a viral epitope, ranging from the immunodominant CMV peptide *NLVPMVATV* to rarer sequences like *FPQSAPHGV* and *CTELKSLDY,* and cell values are five-fold average AUROCs. DeepPROTECTNeo (first row in the heatmap) maintains remarkably consistent performance, with AUROCs spanning only 0.53 to 0.74 and exceeding 0.60 on the majority of under-represented epitopes. By contrast, competing methods such as ERGO-AE and ATM-TCR exhibit greater volatility on frequent epitopes (e.g., ATM-TCR hits 0.78 on *NLVPMVATV*) but often fall below 0.55 on low-prevalence epitopes. Even NetTCR-2 and epiTCR, while being moderately robust, show wider dips around 0.48-0.52 for several targets. Together, these reveal DeepPROTECTNeo’s unique ability to capture both the fine-grained biophysical dependencies of TCR-epitope contacts and the broad, epitope-agnostic patterns required for real-world immunological prediction.

### 2.5 A Case-Study on Patient-specific Clinically Validated Cancer Cohort

To evaluate DeepPROTECTNeo’s clinical relevance, we applied it to five TESLA subjects-three with melanoma (P1, P2, P3) and two with NSCLC (P12, P16), each containing matched RNA-Seq and validated neoepitope data from the TESLA consortium[53]. After filtering out fusions and splicing events due to insufficient validation evidence, our analytical pipeline effectively identified 350 candidates out of the 532 potential peptide–MHC complexes evaluated by TESLA using NetMHCpan-based filtering based on SNV/indels only. Among these 350 candidates, 24 were among the 34 experimentally validated immunogenic epitopes in the TESLA dataset. We further analysed these 350 candidates by pairing each epitope with RNA-Seq-derived TCR sequences from the same patient for TCR binding (detailed patient-wise distribution of detected, validated, and non-validated epitopes is given in Additional File 2, Figure S5). We obtained binding scores for all possible TCR-epitope combinations, and each epitope was summarized using the 95th percentile of its TCR binding scores to capture strong interaction signals while minimizing the impact of outliers. A TCR-specificity criterion of 0.5 was applied to the 95th percentile scores, resulting in the retention of 252 peptide candidates as high-confidence neoepitopes (See score aggregation details in Additional File 1, Note S4). The core discriminative principle of DeepPROTECTNeo, in which retained epitopes consistently exhibit higher 95th percentile TCR scores than non-retained ones (Fig. 7A). This separation was statistically profound (Mann-Whitney U=24696, p=8.1×10^-48^, Cliff’s δ=1.0), demonstrating complete non-overlap of score distributions and substantiating the choice of the 0.5 threshold for epitope retention.

**Figure 7.**
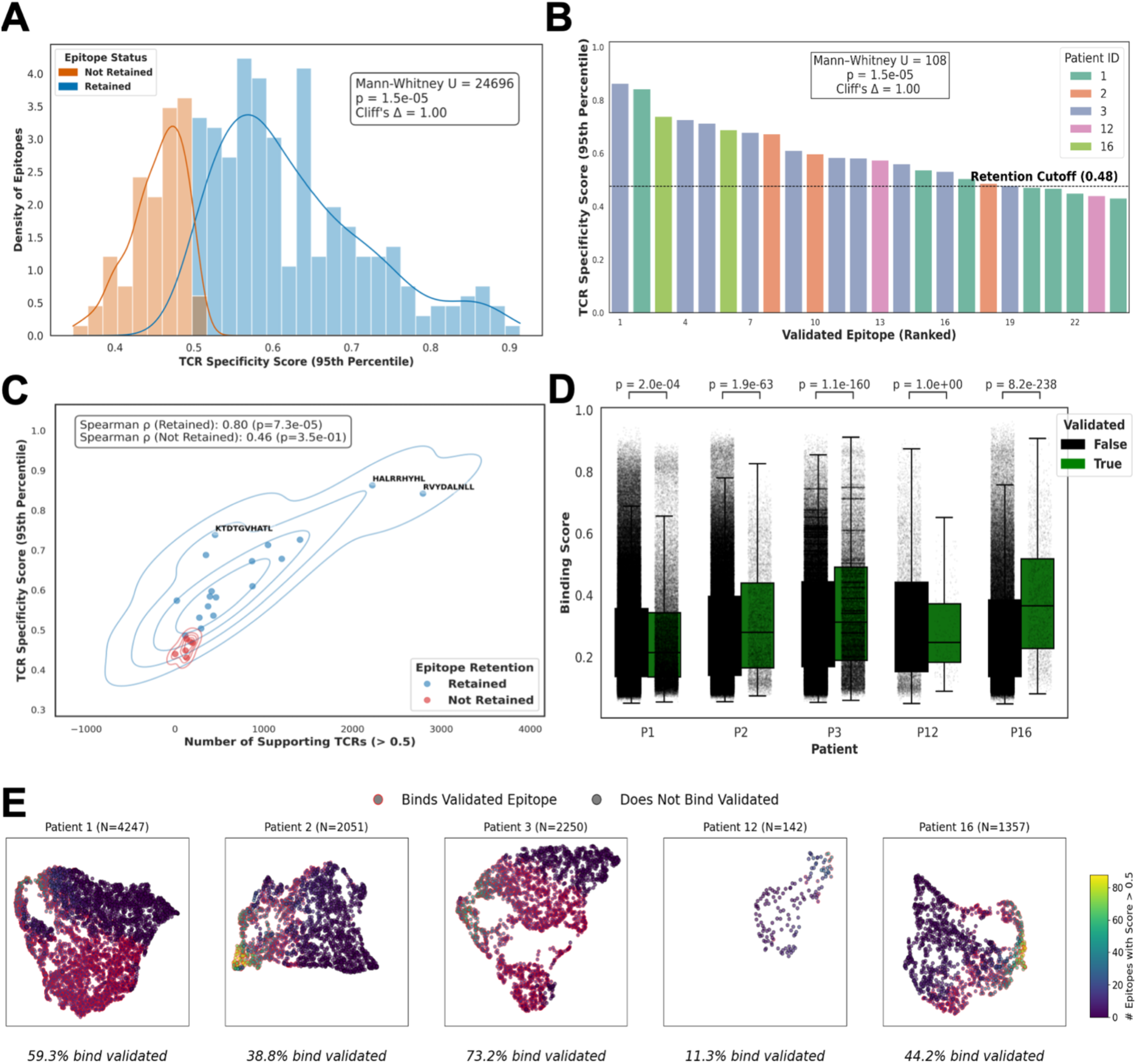
DeepPROTECTNeo captures immunogenic epitopes in TESLA patients via TCR-aware prediction. **A)** Distribution of DeepPROTECTNeo-predicted TCR specificity scores (95th percentile per epitope) for validated (blue) and non-validated (orange) neoepitopes across five TESLA patients. Validated epitopes displayed significantly higher scores (Mann– Whitney *U* = 24,696, *p* = 8.1×10⁻⁴⁸, Cliff’s δ=1.00), supporting strong immunogenic signal capture. **B)** Ranking of 24 validated epitopes by TCR specificity score, stratified by patient. A threshold of 0.48 (dashed line) retained 18 of 24 epitopes (70.83% recall) with a large effect size (Cliff’s δ= 1.00). **C)** Association between epitope TCR specificity and the number of strongly interacting TCRs (*score > 0.5*) for 24 validated epitopes. A strong positive Spearman correlation was observed for retained epitopes (ρ = 0.80, *p* = 7.3×10⁻⁵), but not for non-retained ones (ρ = 0.46, *p* = 0.35), indicating that immunogenic peptides tend to recruit larger numbers of high-affinity TCRs. Contour density overlays show distinct spatial clustering patterns between retained (blue) and non-retained (red) epitopes. The top three epitopes by TCR specificity score: KTDTGVHATL, HALRRHYHL, and RYVDALNLL are annotated, illustrating the immunological prominence of these sequences within the cohort. **D)** Per-patient comparison of DeepPROTECTNeo binding scores between validated and non-validated epitopes. Validated epitopes consistently exhibited significantly higher scores across four patients (Mann–Whitney U test; *p* < 2×10⁻⁴; Bonferroni-corrected), with P3 (*p* = 1.1×10⁻¹⁶⁰) and P16 (*p* = 8.2×10⁻²³⁸) showing the strongest effect. **E)** UMAP projections of patient-specific TCRs coloured by the number of epitopes bound with high affinity (*score > 0.5*). TCRs binding at least one validated epitope (red edges) formed enriched spatial clusters, particularly in P1 and P3, with validated binding rates ranging from 11.3% (P12) to 73.2% (P3).

Further, we demonstrated DeepPROTECTNeo’s 95th-percentile TCR specificity scores for 24 experimentally validated neoepitopes across five patients (Fig. 7B), ranging from 0.431 to 0.863 (median 0.578). For retaining peptides having high TCR specificity, we determined the cutoff score in two steps. First, for each epitope we summarized its predicted TCR-epitope interactions by taking the 95th percentile of all binding scores (“TCR_95P_Score”), followed by collecting those epitopes with TCR_95P_Score≤0.50 and computed the 95th percentile of that subset, yielding a final retention cutoff of 0.48 (Additional File 1, Note S5; Additional File 2, Figure S6). Performing in depth analysis for identifying stage wise elimination of validated epitopes reveals that the upstream MHC-binding filter accounts for the larger share of attrition: 10 of the 34 validated epitopes are excluded as non-SNV/indel variants or sub-threshold HLA-binders, while the TCR-specificity filter retains 18 of the 24 MHC-passing candidates (75 % in-pool recovery; See details about all the epitopes detected in different stages of pipeline in Additional File 3, Table S7, S8, S9; Additional File 2, Figure S7). Two systematic patterns account for nearly all TCR-stage non-retention: four are HLA-A*02:01 FL-motif epitopes from patient 1 that exhibit broad, flat CDR3β score distributions despite this patient’s 338,720-pair repertoire, and one (Patient 12, ALYFNSQWK) reflects extreme repertoire sparsity, only 142 unique CDR3β sequences were recovered from TIL, compared with 1,357–4,247 for the other patients. Per-patient AUC analysis (Additional File 3, Table S10; Additional File 2, Figure S8) reveals strong discrimination where TCR data is adequate (Patient 16: AUC = 0.885, permutation p = 0.032), while Patient 12’s sparse TIL limits discrimination. The retained versus non-retained distributions were perfectly separated (Mann–Whitney U=108, p=1.5×10^-5^, Cliff’s δ=1.00), confirming maximal discrimination at this cutoff. In summary, DeepPROTECTNeo calls 252 of 532 peptides, capturing 18/34 true neoepitopes (52.9% sensitivity) and rejecting 264/498 negatives (53.0% specificity), for a BA of 53.0% and ROC AUC of 0.526. The entire pipeline of DeepPROTECTNeo outputs MHC-epitope binding followed by TCR specificity ranking of the neoepitopes. On the other hand, existing neoantigen predictors TSNAD v2.0[54] and DeepNeo[55] operate only up to the MHC-binding stage. Therefore, we report DeepPROTECTNeo’s matched-scope for MHC-only sub-output as well as its full end-to-end output. Analysing DeepPROTECTNeo against TSNAD v2.0, and DeepNeo on the same 5-patient TESLA cohort[56], we report 24 of 34 (70.59%) matched MHC-only validated epitopes (70.6 % sensitivity, 34.5 % specificity, 52.6 % BA) compared to 11 out of 34 (32.35%) for DeepNeo[55] (sensitivity: 32.4%, specificity: 87.2%, BA: 59.8%, AUC: 0.556) and only 4 out of 34 (11.76%) for TSNAD v2.0[54] (sensitivity: 11.8%, specificity: 96%, BA: 53.9%, AUC: 0.487). Moving further with the additionally filters, the 24 MHC-passing candidates through patient-specific TCR repertoire retains 18 of 24 (75% in-pool recovery), resulting 18 validated neoepitopes from the full end-to-end pipeline (52.9 % sensitivity, 53.0 % specificity, 53.0 % BA, AUC 0.526; Fisher’s exact OR = 8.44 vs. TSNAD v2.0, p = 0.0006; OR = 2.35 vs. DeepNeo, p = 0.141; Additional File 3, Table S11). Cross-referencing these 18 retained epitopes against the five TESLA validation assays (TCR_FLOW_I / II binary multimer, TCR_FLOW_I / II_QUANT quantitative T-cell frequency, and TCR_NANOPARTICLE multimer)[53] confirms that every retained epitope is positive in at least one independent assay, with three high-confidence targets (ATYKGVPYEVK, AINRPTVLK, FTNESYLELY in Patient 3) corroborated across both flow rounds (Additional File 3, Table S12). This matched-scope comparison along with confirmed immunogenicity in different assays establishes that, DeepPROTECTNeo’s end-to-end capability of neoepitope detection is substantially more (38.24% more than DeepNeo and 58.82% more than TSNAD v2.0) sensitive than prior MHC-only tools, and that the TCR-aware stage functions as a principled specificity filter rather than as a source of additional candidates.

Further, characterising DeepPROTECTNeo as a ranking tool rather than a binary classifier, we computed per-patient Precision@K, Recall@K, and fold-enrichment metrics on the 24 MHC-passing validated epitopes (Additional File 3, Table S13; Additional File 2, Figure S9). In Patient 3, the top-ranked epitope (HALRRHYHL) was experimentally validated and 3 of the top 6 candidates were confirmed immunogenic (Top-5 Precision = 40%, 2.24-fold enrichment over baseline). In Patient 16, both validated epitopes appeared within the top 15 of 89 candidates, achieving 100% recall at 17% list depth (5.93-fold enrichment; hypergeometric p = 0.027). Performance correlated with TCR repertoire depth: Patient 12 (TIL-derived; 142 unique CDR3β) showed no enrichment, whereas PBMC-derived patients with deeper repertoires (P1: 338,720; P3: 196,785 TCR–epitope pairs) consistently placed validated epitopes in the upper ranks.

To investigate the statistical significance of these binding scores, we identified retained epitopes that show a strong positive correlation between the number of supporting TCRs and their 95th-percentile score (ρ=0.8, p=7.3×10⁻^5^) (Fig. 7C), whereas non-retained peptides exhibit no such relationship, reinforcing that true immunogenicity arises from convergent, high-affinity TCR engagement rather than stochastic binding. Further, we examined per-patient score distributions (Fig. 7D), showing that validated epitopes consistently receive higher DeepPROTECTNeo scores in 4 out of 5 patients. Notably, P2 (p=1.9×10⁻⁶³), P3 (p=1.1×10⁻¹⁶⁰), and P16 (p=8.2×10⁻²³⁸) showed extremely significant differences, and P1 reached moderate significance (p=2.0×10⁻⁴).

To assess how DeepPROTECTNeo’s 95th percentile TCR binding scores (TCR_95P_Score) complement standard neoantigen features, we trained logistic regression (LR) and random forest (RF) models incorporating features such as MHC affinity, stability, hydrophobicity, foreignness, and the TCR_95P_Score. LR significantly outperformed RF (AUC=0.81 vs. 0.72). Crucially, TCR_95P_Score had the highest odds ratio (∼1.50), surpassing foreignness, agreptopicity, and MHC-binding features, indicating it is the strongest independent predictor of epitope validation, supporting the orthogonality and additive value of TCR-aware information in neoantigen prediction (Additional File 2, Figure S10). UMAP projections of TCR–epitope interactions reveal distinct TCR clusters (Fig. 7E), particularly in P1, where five major clusters were observed (largest: 418 TCRs), indicating clonal expansion. TCRs binding validated epitopes showed significantly lower CDR3β entropy in 4 of 5 patients, P1 (3.25 vs. 3.03, p=4.0×10⁻¹²³), P2 (3.28 vs. 3.14, p=1.6×10⁻¹⁸), P3 (3.25 vs. 3.03, p=6.7×10⁻⁶⁰), and P16 (3.24 vs. 3.04, p=1.7×10⁻²⁹), suggesting repertoire convergence toward immunogenic epitopes. Only P12 showed no significant difference (*p* = 0.49). These patterns validate DeepPROTECTNeo’s capacity to capture immune selection dynamics. Together, these comprehensive results suggest DeepPROTECTNeo significantly enhances both the sensitivity and specificity of immunogenic epitope prioritization, suggesting a critical advancement for designing effective cancer vaccines.

### 2.6 DeepPROTECTNeo Reveals Descriptor Hierarchies and Structural Contacts of TCR–epitope Binding

Using our model’s interpretability, we investigate a clustered heatmap of normalized importance ranks for 82 common peptide descriptors across ImmuneCODE, Mutation, and Viral test sets (See Fig. 8A). Applying agglomerative hierarchical clustering with Euclidean distance and the average-linkage criterion to attention-derived ranks (Additional File 1, Note S6), reveals three clear descriptor families. Across all three cohorts, DeepPROTECTNeo’s attention scores consistently captured 14 descriptors (PRIN1/2, SVGER7/8, ProtFP2/3, AF3/5) to normalized importance >0.8, reflecting their universal role in capturing global physicochemical gradients[57], mid-range sequence-order couplings[58], and short-motif signatures. Conversely, the Mutation dataset specifically engaged specialized descriptors such as BLOSUM7, T3, VSTPV1, and ST3 with moderate significance (0.5–0.7), aligning with their responsiveness to individual amino-acid alterations at canonical anchor sites[59,60]. Similarly, viral epitopes within the Viral set demonstrated an enrichment of hydrophobic and aromatic-frequency characteristics (VHSE4, ProtFP5, VHSE8, ProtFP7), suggesting that these attributes encapsulate conserved glycoprotein motifs[61] critical for the identification of CMV/EBV/Influenza. At the lower end, electronic Z-scale descriptors (E1–E4), 3D-shape WHIM features (MSWHIM3), and low-order SVGER terms (SV2/3) consistently recorded below 0.1 across all datasets, indicating that detailed electronic or inferred tertiary-structure proxies provide negligible predictive value in a sequence-only context[57,62]. This negligible contribution of E1–E4 has two molecular-geometric explanations. Firstly, strong collinearity (|Pearson r| > 0.8) between E1–E4 and existing polarity/electrostatic descriptors in the physicochemical feature set, making them informationally redundant; and next the dominance of steric shape complementarity and short-range electrostatic interactions over long-range electronic-topological effects in CDR3β–epitope binding at the 8–20-residue scale. Cross-cancer-type generalisability is not affected, because the conserved substitution patterns are captured by the BLOSUM62 component of the skip pathway, and the remaining physicochemical features (hydrophobicity, flexibility, accessibility proxies) adequately represent the binding-relevant structural context. A strong pairwise Spearman correlations between ImmuneCODE, Mutation, and Viral datasets (ρ = 0.90-0.94, p<10⁻³⁸) indicates a common core of key descriptors with only a few dataset-specific outliers (Additional File 2, Figure S11).

**Figure 8.**
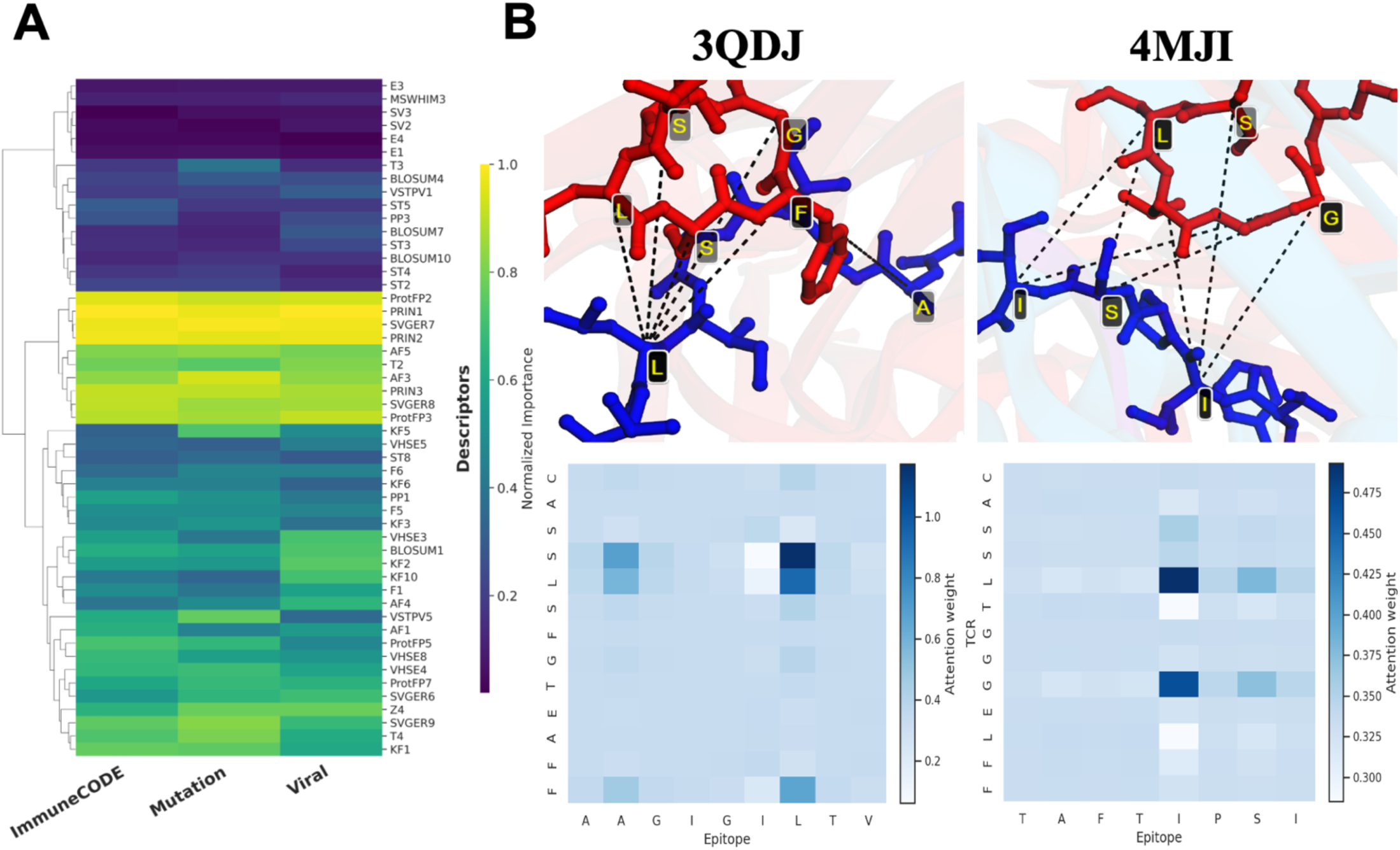
Cross-dataset descriptor prioritization and structural attention mapping. **A)** Non-zero–variance descriptors from ImmuneCODE, Mutation, and Viral sets were ranked by attention importance, merged, normalized, and clustered (Ward’s linkage). The bright yellow block highlights 14 core features (including PRIN1/2, SVGER7/8, ProtFP2/3, AF3/5) consistently prioritized across all datasets; mid-green bands denote dataset-specific specialists; deep-purple tails are low-variance or pruned descriptors. **B)** Top: Two representative TCR– peptide cocrystal structures (PDB 3QDJ, 4MJI) with peptide side chains colored by normalized attention (blue: epitope, red: TCR) and dashed lines indicating sub-8 Å contacts. Bottom: Corresponding sequence-level attention heatmaps, recapitulating the structural hotspots captured by DeepPROTECTNeo.

Further, we mapped DeepPROTECTNeo’s epitope-side attention onto two representative TCR–epitope cocrystal structures (PDB ID: 3QDJ, 4MJI; Fig. 8B). In 3QDJ (left panel), five sub-8Å contacts: A2–F97 (7.40Å), L7–S94 (7.11Å), L7–L95 (3.55Å), L7–S96 (3.62Å) and L7–F97 (5.81Å) colocalize with the highest attention at P2 and P7. Distances of 3.5-3.6Å between L7 and L95/S96 reflect tight van der Waals packing and hydrogen bonding in the CDR3 loop[61], while the 5-7Å separation to F97 denotes stabilizing aromatic edge-to-face interactions[63]. Similarly, in 4MJI (right panel), four contacts under 8Å: I5–G98 (7.58Å), S7– L95 (3.62Å), S7–G98 (7.98Å), and I8–L95 (4.33Å), align with pronounced attention at mid-peptide positions, underscoring the model’s sensitivity to sequence-order couplings and aromatic-frequency signals[64]. By recovering these sub-8Å hydrogen bonds, van der Waals contacts, and aromatic stacking purely from sequence attention, DeepPROTECTNeo captures the very molecular grammar that underlies TCR-peptide specificity and, by extension, the key features leveraged in reverse-vaccinology epitope selection[65]. Together, the cross-dataset heatmap (Fig. 8A) and structural projections (Fig. 8B) demonstrate that DeepPROTECTNeo’s sequence-only attention mechanism distils a minimal, biologically meaningful set of descriptors using interpretable feature prioritization with direct recapitulation of key noncovalent contacts, while offering both mechanistic insight into the molecular grammar of TCR recognition and robust predictive generalization across diverse epitope datasets. Additionally, to validate the interpretability of DeepPROTECTNeo’s attention at population scale, we extended the analysis to five independent quantitative benchmarks. First, across 81 TCR–pMHC crystal structures, the model’s residue-pair attention matrix discriminates true ≤ 5 Å physical contacts from non-contacts with a median per-PDB AUROC of 0.636 (one-sample t-test vs. 0.5, p = 8.4 × 10⁻⁸; Additional File 3, Table S14, Additional File 2, Figure S12). Second, a per-PDB permutation baseline (1,000 shuffles) confirms that this signal is not a scoring artefact; the real median AUROC 0.636 significantly exceeds the permuted median 0.500 (paired t-test, p = 1.0 × 10⁻⁷; Additional File 3, Table S15; Additional File 2, Figure S13). Third, restricting to the 34 HLA-A*02:01 9-mer structures, the mean attention on the solvent-exposed TCR-contact zone (P3–P6) is 0.1265, vs. 0.0606 on the MHC-buried anchor residues (P2, P9), a 2.09-fold difference present in 34/34 PDBs (paired t-test, p = 1.9 × 10⁻²⁵; Additional File 3, Table S16; Additional File 2, Figure S14), recovering a textbook TCR– pMHC feature without any positional or structural supervision during training. Fourth, side-by-side attention-vs-contact visualisations for the three highest-AUROC PDBs (5WKH, 5WKF, 6RPA) confirm the signal at the individual-structure level (Additional File 2, Figure S15). Fifth, on the 4,244-pair deep-mutational-scan panel, positional attention is significantly enriched at mutation-critical epitope residues across 26 independent CDR3β clonotypes (Wilcoxon signed-rank, p = 0.024; 16/26 TCRs show ρ > 0; Additional File 3, Table S17; Additional File 2, Figure S16). Together, these five analyses establish that the attention-based interpretability signal shown in Fig. 8 is not anecdotal but systematic and biologically meaningful.

## 3 Discussion

DeepPROTECTNeo is an end-to-end framework for neoantigen prioritization, integrating raw RNA-Seq preprocessing, HLA typing, external MHC-peptide affinity scoring via NetMHCpan, and a unique CDR3β-epitope binding prediction module within a unified pipeline. Two-branch structure of DeepPROTECTNeo combines backbone-constrained knowledge with sequence-derived determinants, helping to learn both long-range dependencies and local spatial correlations that are important for TCR-epitope pairing. Our model yields state-of-the-art predictive accuracy, strong early-retrieval of true binders and minimal inter-fold variability, while offering mechanistic insights that align with decades of structural immunology. DeepPROTECTNeo generalizes robustly across all three benchmarks and exhibits strong quantile concordance with unbiased filtered evaluation (no independent test pair is present in stratified five-fold cross-validation data, see Additional File 3, Table S18), achieving the highest alignment between ImmuneCODE and Viral (Pearson r = 0.99), substantial agreement between ImmuneCODE and Mutation (r = 0.91), and robust, if slightly lower, concordance between Mutation and Viral (r = 0.89) demonstrating both broad generalization and dataset-specific calibration of predictive confidence (Additional File 2, Figure S17). Central to DeepPROTECTNeo is its multi-scale interpretability: attention-guided motif extraction and descriptor-based embeddings converge on a concise set of biochemical signatures like global physicochemical axes (PRIN), sequence-order couplings (SVGER), and short-motif fingerprints (ProtFP), which consistently map onto atomic contacts within 8Å, thereby reconstructing the core TCR-peptide contacts purely from sequence.

However, we acknowledge several limitations of the current framework that point to clear directions for future improvement. DeepPROTECTNeo infers interaction specificity using the CDR3β chain only, matching the dominant single-chain paradigm of existing state-of-the-art methods (ATM-TCR, NetTCR-2.0, ERGO-AE/LSTM, epiTCR, TEINet); because physiological TCR–epitope recognition is governed by paired αβ conformational geometry, β-only inference inherently limits structural oversight. The architecture has nevertheless been designed so that extension to paired αβ data requires only instantiating a parallel α-chain extraction branch (reusing the BiLSTMEncoder + DeepCNNAttnProjector design) whose output concatenates alongside the β-chain vector prior to cross-attention — an extension contingent on wider availability of paired single-cell sequencing. Relatedly, training uses 1:1 random-recombination negatives; although the CD-HIT-85% hard-negative experiment (108,543 pairs) and the 1:2 imbalanced clinical-prevalence experiment (338,382 pairs) in Section 2.1 demonstrate robustness to pseudo-negatives and to class imbalance, fully experimentally-verified non-binding pairs remain rare in public databases and are an intrinsic constraint of the training distribution. With respect to clinical generalisation, the pooled TESLA AUC is depressed by two mechanistically identified sources i.e. FL-motif HLA-A*02:01 epitopes and TIL-derived repertoires with < 200 unique CDR3β limiting generalisation in these specific sub-cohorts; broader evaluation on larger clinical cohorts beyond melanoma and NSCLC will therefore be essential. Continued advances in single-cell sequencing, paired αβ registries, deep structural modelling, and structure-informed representation learning offer promising avenues to address these constraints.

In conclusion, DeepPROTECTNeo represents a substantial advance in computational neoepitope prioritization for immunotherapy by integrating a well-calibrated predictive model with mechanistic interpretability. Its multi-scale fusion and attention-based architecture enables accurate, biologically grounded feature prioritization, distilling a minimal set of descriptors that recapitulate key noncovalent contacts underlying TCR recognition. This combination of domain-informed design, strict cross-dataset benchmarking, and interpretable sequence-only modelling delivers both sensitivity and specificity gains in epitope ranking, supporting more effective cancer vaccine design. Moreover, the architecture provides a flexible foundation for integrating α/β TCR pairing, detailed antigen-processing events, experimentally derived negatives, and validated structural representations, positioning DeepPROTECTNeo as a platform for next-generation immuno-informatics.

## 4 Conclusions

In this article, we introduce DeepPROTECTNeo, a unified, clinically deployable deep learning pipeline for personalized neoantigen prioritization. DeepPROTECTNeo streamlines the entire workflow starting from raw sequencing data through unified postprocessing frameworks to prioritize TCR-epitope binding through a reverse vaccinology-inspired biologically informed architecture. Furthermore, the framework integrates fine-grained residue-level interactions via a rotary cross-attention mechanism, enabling both robust score prediction and highly accurate mechanistic interpretability of key anchor residues and contact motifs. Clinically, its utility is established by successfully separating true binders within the experimental TESLA consortium cohort and additionally capturing critical clinical features, outperforming state-of-the-art methods with consistent high binding scores across the viral and mutational independent datasets.

## 5 Methods

### 5.1 Data Processing

DeepPROTECTNeo begins with processing of RNA-Seq FASTQ files using FastQC[66] for base quality, GC content, and read-length distributions, then adapters and low-quality bases are trimmed using Trimmomatic[67]. High-quality reads are aligned to the GRCh38/hg38 reference genome via HISAT2[68], and the resulting BAM files are processed with SAMtools[69] for filtering unmapped or low-quality reads, sorting, marking duplicates, realigning indels, and recalibrating base qualities to produce high-fidelity alignments. Somatic variants (SNVs and indels) are called using GATK4 Mutect2[70] and annotated with ANNOVAR[71] against refGene[72], ensGene[73], gnomAD[74], ClinVar[75] and COSMIC[76], retaining only exonic, non-synonymous mutations for neoepitope generation. We used germline-resource (gnomAD) and p-o-n (Panel of Normals) to reduce the false positive variants resulting from Mutect2. Patient HLA class I and II alleles are inferred directly from these BAMs using arcasHLA[12], while CDR3β sequences are extracted in parallel using TRUST4[77]. Each candidate neoepitope spanning an identified mutation is then scored for peptide–MHC binding affinity with NetMHCpan-4.1 and NetMHCIIpan-4.0[14] for both MHC (I and II) and high-confidence neoantigens are selected via threshold-based filtering on binding affinity score (≤ 0.5). Further, filtered epitopes are ranked for TCR-binding affinity using DeepPROTECTNeo and NetBCE[78] is used for proving BCR affinity.

### 5.2 TCR-epitope Interaction prediction

DeepPROTECTNeo employs a dual-branch architecture that fuses sequence and physicochemical information for TCR–epitope binding prediction. The token branch embeds CDR3 and peptide sequences into 240-d vectors with learned positional encodings, then applies BiLSTM and attention pooling. Simultaneously, the skip branch processes 102-d descriptors through 32 Conv1D + SE blocks (channel width *C* = 64, kernel size 5) and optional self-attention, pooling intermediate and final maps into a 30-d skip summary. A gated fusion merges each 240-d token feature with its 30-d skip, a 6-head rotary cross-attention captures inter-sequence context, and the concatenated 720-d vector is refined by 6-head self-attention and a 24-layer ReZero RetNet before final MLP classification.

Let *T* = (*t*_1_, …, *t_n_*), *P* = (*p*_1_, …, *p_m_*) be the CDR3β and epitope sequences over the 20 amino acid alphabets. We learn *f*: (*T*, *P*) → {0,1} to predict binding. Internally, *T* and *P* are mapped to continuous representations via embedding, positional encoding, and feature projection, thereby processed through parallel branches, fused and cross-attended, refined by a ReZero network, and finally classified by a two-way MLP.

#### 5.2.1 Token Branch

To convert discrete tokens into continuous representations, we use a shared embedding matrix of dimension *d* = 240, and we add learned positional encodings so the model knows each residue’s absolute position along the sequence. Concretely, for the TCR:

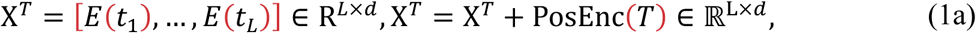

and similarly for the epitope X*^P^* ∈ ℝ*^L^*^×*d*^. *L* can be *m* or *n* based on the length of TCR or epitope. The positional embeddings (PosEnc (T/P) ∈ ℝ*^L^*^×*d*^) are trained jointly with the rest of the network, letting the model learn which positions are most informative. X is fed through a two-layer bidirectional LSTM (hidden size *d*/2 per direction). Denote by

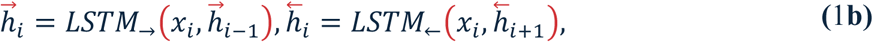

the forward and backward states at position *i*. We concatenate to form 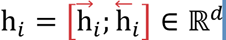 and collect H = (h_1_, …, h*_L_*) ∈ ℝ*^L^*^×*d*^ for P and T.

#### 5.2.2 Skip Branch: DeepCNNAttnProjector and SkipGate

Handcrafted peptide descriptors from both TCR sequences and epitopes *F* ∈ ℝ*^B^*^×102^are first reshaped to *H*^(0)^ = *F*.unsqueeze(1) ∈ ℝ*^B^*^×1×102^. This tensor is passed through a stack of 32 residual Conv1D blocks. At block *i*, the input *H*^(*i*−1)^ undergoes two Conv1D-BatchNorm-GELU steps and is added back to a skip connection:

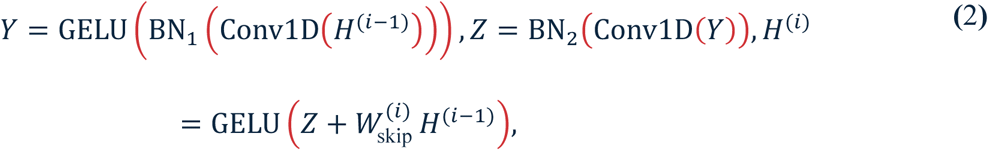

Here 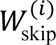 is the identity unless a 1 × 1 projection is needed to match dimensions. After 32 blocks, a 1 × 1 convolution reduces channels to *D* = 60:

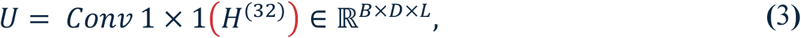

A channel attention module then re-weights these channels by first computing

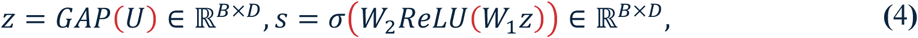

and scaling *U*_CA_ = *U* ⊙ *s*.unsqueeze(−1). A multi-head self-attention layer refines this to *U*_SA_. A final GAP produces the 60-dim feature *t*_final_ = GAP(*U*_SA_) ∈ ℝ*^B^*^×60^.

Each intermediate feature map *H*^(30)^ and *H*^(31)^ is globally averaged and pooled to ℝ*^B^*^×64^, feature map *t*_final_ to ℝ*^B^*^×60^. Each resulting vector is then passed through its linear transformation to produce three 30-dim skip vectors.

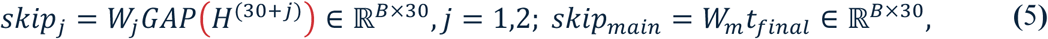

Concatenating these three 30-dim vectors yields *S* ∈ ℝ*^B^*^×90^, followed by a final ReLU-MLP “SkipGate” reduces to the 30-dim summary

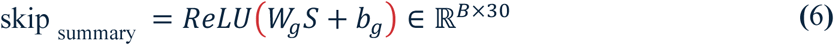

ready for fusion with the token-branch features.

#### 5.2.3 Gated Fusion

Let **F***^t^* ∈ ℝ*^B^*^×*d*^_model_ be the batch of token-branch features and F*^s^* ∈ ℝ*^B^*^×*d*^_skip_ the batch of skip-branch summaries. We first project both into the common *d*_model_-dimensional space:

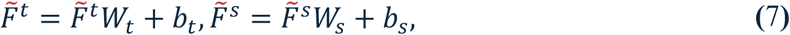

where *W_t_*, *W_s_* ∈ ℝ*^d^*^model ×*d*model^ and *b_t_*, *b_s_* ∈ ℝ*^d^*^model^. Next, we compute an element-wise gate g ∈ (0,1)*^B^*^×*d*model^ from their concatenation:

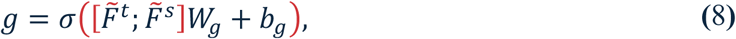

With *W*_@_ ∈ ℝ^2*d*model ×*d*model^ and *b*_@_ ∈ ℝ*^d^*^model^. Finally, the fused feature **F**_fused_ ∈ ℝ*^B^*^×*d*model^ is a convex combination:

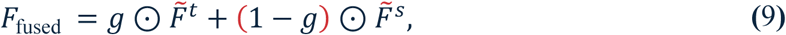

allowing the network to dynamically balance sequence-level and physicochemical information on a per dimension basis.

#### 5.2.4 Rotary Positional Embedding and Cross-Attention

Rotary Positional Embeddings (RoPE) was used to capture relative offsets by rotating each head’s query and key vectors in the complex plane by an angle dependent on token position *p*. Concretely, for a *d_k_*-dimensional vector *x_p_*, we split even and odd indices and rotate:

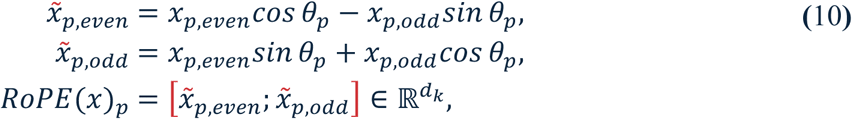

where *θ_p_* is a fixed (or learned) rotation per position. Applying the same RoPE to both queries and keys ensures that their dot-product depends only on the relative distance Δ = *p* − *q*.

Given fused TCR and peptide features 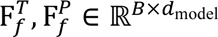, we project into multi-head queries, keys, and values:

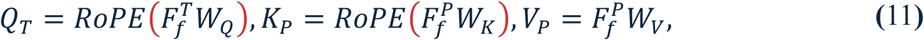

and compute cross-attention in both directions:

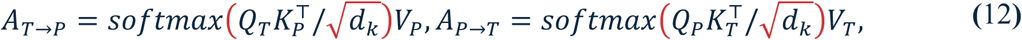

We then symmetrize by averaging:

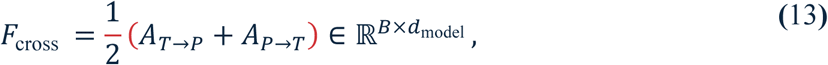

This bidirectional cross-attention captures inter-sequence dependencies and relative positional relationships in a single, unified operation.

#### 5.2.5 RetNet and Self-Attention

Let 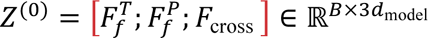, denote the concatenated TCR-fused, peptide-fused, and cross-attended features. This tensor is pro two parallel streams. In the Enhanced RetNet stream, we apply *L* ReZero-style residual layers

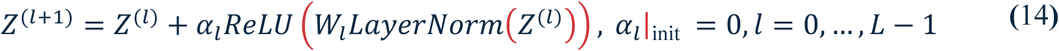

Initializing each *α_l_* = 0 ensures that the network begins as an identity mapping, providing improved gradient flow and training stability, so that after *L* layers we get *F*_retnet_ = *Z*^(*L*)^ ∈ _ℝ_*B*×3*d*model.

Concurrently, the Extra Self-Attention stream applies a single multi-head attention block to the original *Z*^(0)^ to obtain *F*_SA_ = SelfAttnM(*Z*^(0)^) ∈ ℝ*^B^*^×3*d*model^.

The final representation is obtained by element-wise summation of these two pathways:

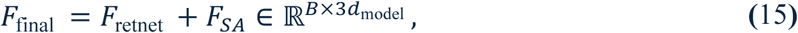

which is then fed into the downstream MLP, followed by a sigmoid for final classification. Ablation of all the major components of our architecture are shown in Additional File 3, Table S4.

### 5.3 Loss

To mitigate over-confidence and tolerate label noise, we train with a label-smoothed cross-entropy. Given a batch of *N* examples with ground-truth labels *y_i_* ∈ {0,1}, the network produces logits 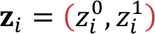 and probabilities

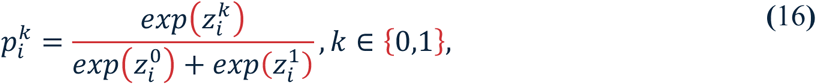

We then define a “soft” target distribution 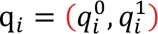 by smoothing the one-hot label:

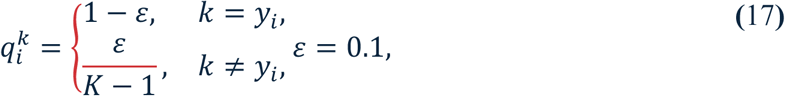

Hence, the final loss averaged over the batch is

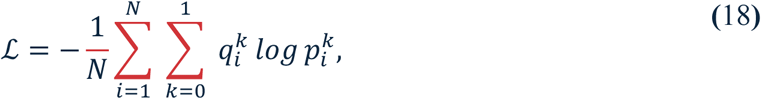

This label smoothing reduces the model’s tendency to become over-confident on the training labels, improving generalization and calibration.

## 6 Declarations

### 6.1 Ethics approval and consent to participate

Not applicable.

### 6.2 Consent for publication

Not applicable.

### 6.3 Data Availability

The raw data were downloaded from publicly available databases (McPAS-TCR[79], VDJdb[80], IEDB[81]). Processed Mutation and Viral datasets used for independent benchmarking was obtained from Zenodo[82] (https://doi.org/10.5281/zenodo.15025579) of ePytope-TCR[43]. For aligning raw sequencing files, we used Human Genome assembly GRCh38 available via HISAT2 Download page (https://daehwankimlab.github.io/hisat2/download/)[68]. We downloaded and installed NetMHCpan-4.1 and NetMHCIIpan-4.0 for MHC-peptide binding, which is available via DTU Health Tech (https://services.healthtech.dtu.dk/services/NetMHCpan-4.1/)[14] and https://services.healthtech.dtu.dk/services/NetMHCIIpan-4.0/). Variant annotation using GATK4 Mutect2 required Homo Sapiens Assembly FASTA for best practices short variant discovery from RNASeq, which was obtained via Google Cloud bucket. For annotating identified variations, all the reference databases were directly downloaded from the ANNOVAR[71] website (https://annovar.openbioinformatics.org/en/latest/user-guide/download/). 3D crystal structure of TCR-MHC-peptide complex (PDB ID: 3QDJ and 4MJI) were downloaded PDB database (https://www.rcsb.org/). For clinical validation of our model, we used the TESLA cohort[53], which is available via Synapse (https://www.synapse.org/) with ID syn23446508. The complete DeepPROTECTNeo workflow is available as a web server via https://cosmos.iitkgp.ac.in/DeepPROTECTNeo/. The source code is available via GitHub (https://github.com/debraj-55555/DeepPROTECTNeo) under the MIT License and all the processed data used in this study for model building, validations, and trained model weights are available via Zenodo[83] (https://doi.org/10.5281/zenodo.16751524).

### 6.6 Author contributions

D.D. and S.B. conceived the study. D.D., S.B, and P.M. contributed to the conceptualization and methodology. D.D. and S.B. contributed to data curation, model implementation, validation, and analysis with baselines, and website development. D.D., along with S.B., conducted the validation on clinical data and neo-epitope extraction methodology design. P.M. D.D., S.B. wrote the manuscript and prepared figures. P.M. provided supervision and contributed to the interpretation of predictive results. All authors contributed to the original draft and critically revised the manuscript. All authors read and approved the final manuscript.

## 6.7 Acknowledgements

Authors are thankful to Avik Pramanick for useful discussion.

## Additional files

### Additional file 1. Supplementary Notes S1-S6

**Note S1.** Data Curation and Pre-processing

**Note S2.** Model Training and Evaluation

**Note S3.** Performance Evaluation Metrics

**Note S4.** TCR-Epitope Binding Score Aggregation

**Note S5.** Retention Threshold Calculation

**Note S6.** Hierarchical Clustering of Cross-Dataset Feature Importance

### Additional file 2. Supplementary Figures S1–S17

**Fig S1.** Overview of the datasets used and their distribution.

**Fig S2.** Overview of the performance across the epochs.

**Fig S3.** Shape-annotated forward-pass block diagram of DeepPROTECTNeo.

**Fig S4.** The effect of the feature projector components on our architecture.

**Fig S5.** Per-patient comparison of epitope counts from the TESLA consortium versus our MHC- and TCR-binding predictions.

**Fig S6.** Illustrating the distribution of TCR specificity scores (95th percentile) across retained and non-retained epitopes.

**Fig S7.** Missed epitope analysis for the TESLA cohort.

**Fig S8.** Per-patient AUC analysis.

**Fig S9.** Per-patient ranking performance on the TESLA cohort: cumulative recall and Top-K fold-enrichment of validated epitopes.

**Fig S10.** Integrating TCR Context Substantially Improves Neoantigen Prioritization.

**Fig S11.** Scatter plots showing pairwise comparisons of peptide-descriptor importance ranks between (left) ImmuneCODE vs. Mutation, (middle) ImmuneCODE vs. Viral, and (right) Mutation vs. Viral.

**Fig S12.** Structural validation of attention-based contact prediction across 81 TCR–pMHC crystal structures (per-PDB AUROC histogram; AUROC vs. contact density).

**Fig S13.** Real vs. permuted AUROC distributions across 81 PDBs. Real median 0.636 significantly exceeds permuted median 0.500 (paired t-test p = 1.0 × 10⁻⁷).

**Fig S14.** Per-position mean attention (P1–P9) across 34 HLA-A*02:01 9-mer PDBs (±95% CI). Red = TCR-contact zone (P3–P6); grey = MHC-buried anchors (P2, P9). Mean: 0.1265 vs. 0.0606 (2.09-fold; p = 1.9 × 10⁻²⁵).

**Fig S15.** Side-by-side attention (a) and contact (b) visualisations for the three highest-AUROC PDBs (5WKH, 5WKF, 6RPA). Left = attention heatmap A[i,j]; right = experimental contact map (≤5 Å).

**Fig S16.** Per-TCR attention vs. mutation-sensitivity scatter across 26 CDR3β clonotypes. Spearman ρ annotated per panel. Blue = ρ > 0; grey = ρ < 0. Wilcoxon p = 0.024.

**Fig S17.** Quantile–Quantile Comparison of DeepPROTECTNeo Predictions Across Independent Datasets.

### Additional file 3. Supplementary Tables S1–S18

**Table S1.** Positive unique pairs from each dataset.

**Table S2.** Module-by-module summary of the DeepPROTECTNeo architecture (12 modules; total 4,921,338 parameters; 19.52 MB FP32). B = batch size; L = sequence length (L_TCR = 20, L_pep = 11); d_model = 240.

**Table S3.** The range of hyperparameters used during the grid search and the best value achieved across the range of value used during training and optimization of our model.

**Table S4.** 9-configuration 5-fold ablation study. Values = mean ± SD across 5 folds. p-values from paired two-sided t-tests vs. C0. * p<0.05; ** p<0.01; *** p<0.001.

**Table S5.** Performance on Viral data.

**Table S6.** Performance on Mutation data.

**Table S7.** Stage-by-stage attrition through the DeepPROTECTNeo pipeline for all 34 TESLA validated epitopes. Status: Retained = passes TCR filter at 0.5; MHC-missed = excluded at MHC stage; TCR-missed = in MHC pool but TCR_95P_Score < 0.484.

**Table S8.** Characterization of the 6 TCR-stage missed epitopes (TCR_95P_Score ≤ 0.484; in MHC pool). Score = TCR_95P_Score (95th percentile of all CDR3β–epitope binding scores for that patient). Gap_to_threshold = 0.484 − Score. Total_TCR_pairs_patient = number of CDR3β–epitope binding pairs scored for the patient. TCRs_above_0.5 = supporting TCRs for that epitope.

**Table S9.** Characterization of 10 TESLA validated epitopes excluded at the MHC-binding filter stage. All 10 are excluded due to sub-threshold predicted HLA binding (%rank > 0.5) or non-SNV/indel variant origin.

**Table S10.** Per-patient AUC with bootstrap 95% CI and permutation p-values. Per_patient_AUC is computed from TCR_95P_Scores of validated vs. all non-validated candidates. Permutation_p: fraction of 10,000 permutations achieving AUC ≥ observed. Patient 16: AUC = 0.885, p = 0.032.

**Table S11.** Fisher’s exact test pairwise comparisons for recovery of 34 TESLA validated epitopes. OR = odds ratio. DeepPROTECTNeo vs. TSNAD v2.0: OR = 8.44, p = 0.0006.

**Table S12.** TESLA assay-level validation of the 18 neoepitopes retained end-to-end by DeepPROTECTNeo at the 0.484 cutoff. Five orthogonal T-cell assays from the TESLA consortium: TCR_FLOW_I and TCR_FLOW_II are independent rounds of binary multimer-based flow cytometry; TCR_FLOW_I_QUANT and TCR_FLOW_II_QUANT report quantitative reactive-T-cell frequency (% of CD8⁺ T cells); TCR_NANOPARTICLE is a nanoparticle-based pMHC multimer assay. ‘+’ = positive; ‘−’ = negative or not reported. All 18 retained epitopes are positive in at least one assay; three high-confidence Patient-3 targets (ATYKGVPYEVK, AINRPTVLK, FTNESYLELY) are corroborated across both flow rounds.

**Table S13.** Per-patient Top-K metrics (K = 5, 10, 15, 20, 30). Fold_Enrichment = Precision@K / Baseline_rate; Hypergeom_p = hypergeometric p-value for observing ≥ TP in top-K.

**Table S14.** Structural interpretability validation across 81 TCR–pMHC crystal structures. contact_auroc = per-PDB AUROC for discriminating true ≤5 Å contacts from non-contacts using A[i,j]. Median contact_auroc = 0.636; one-sample t-test vs. 0.5: p = 8.4 × 10⁻⁸.

**Table S15.** Per-PDB permutation baseline. real_auroc vs. perm_median: paired t-test p = 1.0 × 10⁻⁷. perm_p_per_pdb = fraction of 1,000 shuffles with AUROC ≥ real.

**Table S16.** Per-position attention weights on 34 HLA-A*02:01 9-mer crystal structures. anchor_mean_P2P9 = mean attention at MHC-buried anchor (P2, P9); nonanchor_mean_P3P8 = TCR-contact zone (P3–P8). anchor_wins = 0 for all 34 PDBs. Paired t-test p = 1.9 × 10⁻²⁵.

**Table S17.** Per-TCR Spearman correlation between positional attention weights and mutation sensitivity (4,244-pair deep-mutational scan, 26 CDR3β clonotypes). rho_attn_vs_sens > 0 in 16/26 TCRs; Wilcoxon signed-rank p = 0.024.

**Table S18.** Consolidated summary of all five datasets used in DeepPROTECTNeo training and evaluation. Source column omitted for brevity; see Supplementary S1 for full source citations.

