## Additional File 1 for "DeepPROTECTNeo: A Context-aware Personalized and Reverse Vaccinology-guided Deep Learning Framework for Immunogenicity Prediction"

**(Additional File 1: Supplementary Notes S1-S6)**

Pralay Mitra, Ph.D.

Department of Computer Science and Engineering,

Indian Institute of Technology Kharagpur

West Bengal - 721302, India

ORCID ID: 0000-0003-4119-3788

Note S1. Data Curation and Pre-processing

The unified dataset was constructed by integrating four major TCR-epitope pairing resources, (1) McPAS-TCR^1^: A manually curated database of pathology-associated T-cell receptor sequences, providing a broad range of disease-relevant TCR-epitope pairs, (2) VDJdb^2^: A curated database of TCR sequences with known antigen specificities, emphasizing experimentally validated TCR-epitope interactions, (3) IEDB Class I and II^3^: The Immune Epitope Database (IEDB) offers TCR-epitope pairs for both Class I and Class II MHC restrictions, with Class I focusing on cytotoxic T-cell responses and Class II on helper T-cell responses. All datasets were pre-processed to a unified format, ensuring consistent representation of CDR3 sequences, epitope sequences, MHC restriction, and source annotations. Redundant TCR-epitope pairs present in multiple databases were identified and removed to prevent data leakage and over-representation. Epitope sequences were standardized (e.g., trimming, case normalization) and mapped to a unified nomenclature to avoid fragmentation of identical or highly similar epitopes across datasets. To train robust models capable of distinguishing true TCR-epitope interactions from random pairings, a negative sampling approach was employed. For each positive TCR-epitope pair, negative samples were generated by randomly pairing the same TCR with unrelated epitopes and vice versa, ensuring that negative pairs did not overlap with any known positives in the combined dataset. Care was taken to avoid introducing bias by matching the distribution of negatives to that of the positives in terms of epitope length and CDR3β composition. The dataset was partitioned such that no CDR3β sequence in the validation or test sets appeared in the training set, ensuring that model evaluation reflected true generalization to unseen TCRs. The split was performed after epitope unification, guaranteeing that both TCR and epitope splits respected the final, harmonized epitope definitions. McPAS contributed the largest fraction of unique CDR3s (58.3%) and a substantial portion of unique epitopes (40.9%), reflecting its broad coverage of disease-associated TCRs. VDJDB provided the greatest diversity of epitopes (55.4%), complementing McPAS with experimentally validated pairs. IEDB Class I and II contributed a smaller but valuable set of pairs, particularly for benchmarking model performance on less-represented MHC contexts. The final combined dataset comprised 18,296 unique CDR3 sequences, 735 unique epitopes, and 20,729 TCR-epitope pairs, representing a comprehensive and rigorously curated resource for TCR specificity modelling. This integrated and meticulously curated dataset ensures high-quality, non-redundant, and biologically diverse training data. The unified epitope strategy, strict TCR split, and robust negative sampling collectively enable fair benchmarking and realistic assessment of model performance on novel TCR-epitope interactions. The McPAS-TCR database, we selected entries containing both CDR3β amino acid sequences and epitope peptides, excluding sequences with invalid characters (e.g., asterisks) and retaining only those annotated with MHC class I (MHCI) molecules. This yielded 10,361 unique CDR3β-epitope pairs. VDJdb entries were filtered to include only TCRβ (TRB gene) sequences with a positive confidence score (Score > 0). After removing duplicates, 5,902 high-confidence pairs were retained. The IEDB dataset, comprising both MHC class I and II interactions, required additional processing. We extracted relevant columns containing epitope peptides and curated CDR3β sequences, removed any entries with ambiguous characters or biologically irrelevant annotations (e.g., beryllium atoms), and excluded epitopes longer than 17 amino acids. This step resulted in a large pool of 159,956 CDR3β-epitope pairs. All three datasets were then concatenated into a unified data frame, where we retained only MHC-restricted interactions and further removed duplicate CDR3β-epitope pairs across datasets to prevent bias due to redundancy. The final integrated dataset consisted of 140,992 unique CDR3β-epitope pairs. This dataset represents a comprehensive and non-redundant collection of experimentally validated TCR-antigen interactions, which serves as the foundation for subsequent computational modeling and epitope prediction tasks.

Note S2. Model Training and Evaluation

We set up a strict training and evaluation protocol for DeepPROTECTNeo to make sure that its TCR–epitope binding predictions module is robust and generalizable in settings where the throughput is noisy and low. We used a strict TCR-based stratified five-fold cross-validation strategy to build the model. Each fold only had unique CDR3β sequences. We have taken special care to select those pairs in the test and validations sets and used a strict TCR split criteria in *sk-learn’s ‘GroupShuffleSplit’* (or group by TCR, strict TCR split) during preparing our test, validation and also during hyperparameter optimization.

We trained the model on a dedicated server with an Intel i9 12th Gen processor, 64 GB of RAM, and an NVIDIA RTX A5000 GPU with 24 GB of VRAM. We provide a unified dataset (140,992 unique CDR3β-epitope pairs among 130,303 TCRs and 1,903 epitopes) for further training and validation of our model in the GitHub provided along with the code. We used the *PyTorch* deep learning framework for training. GPU acceleration was necessary to deal with the multi-branch model's large number of parameters and input batch sizes in a reasonable amount of time. Systematic negative sampling was used to add to the training data at each fold to make sure there were equal numbers of positive and negative pairs. All TCR and epitope sequences were pre-processed to a standard format, with noncanonical characters filtered out and length limits set, as explained in Supplementary S1. We initialized and optimized a deep copy of the model from scratch for each fold using the *AdamW* optimizer (learning rate 0.0005). This allowed for independent convergence and unbiased performance estimation. To avoid overfitting, we used early stopping based on validation loss. We also used the minimum validation loss as a critical checkpoint in zero shot settings to choose the best model. We used a large grid search to optimize the hyperparameters, looking at important architectural parameters like cnn_blocks, cnn_dim, skip_dim, d_model, n_heads, and so on. We then used ANOVA F-scores and validation loss to measure performance. The best-performing setup had a model dimension (d_model) of 240, 3 transformer heads, 32 deep CNN blocks, 64 CNN channels, a kernel size of 5, and a 36-dimensional skip connection pathway. These values were very close to the best ones found in Supplementary S4 and Table 2. The whole model was trained from start to finish for each fold. It included bidirectional LSTM encoding branches, convolutional feature extractors for physiochemical descriptors, gated fusion modules, rotary cross-attention, and multi-scale skip connections. We calculated all of the training and validation metrics, such as balanced accuracy, F1, AUROC, and AUPRC (see Supplementary S3 for more information), both per fold and as the mean±s.d. over the five splits. This made sure that the results were significant and could be repeated. We retrained competing models (ATM-TCR^4^, ERGO-AE^5^, ERGO-LSTM^5^, TEINet^6^, epiTCR^7^, and NetTCR-2.0^8^) from scratch on the same splits, using their default or recommended settings. This made sure that DeepPROTECTNeo's input and output requirements were met for a fair comparison. We used SoftMax activation to process the model outputs and get probability scores. We calculated all of the evaluation metrics as described in Supplementary S2.
DeepPROTECTNeo is also used as a web server, using the same model weights and code as the research workflow. Users who don't have access to high-performance computing can use the model through an easy-to-use web interface (<https://cosmos.iitkgp.ac.in/DeepPROTECTNeo/>). This lets them do full analysis on raw FASTQ/BAM files or quick evaluations on pre-paired CDR3-epitope inputs. There is example jobs based on external clinical data, like TESLA Patient 1. The web system runs on the same hardware as the research training (A5000 GPU, 64GB RAM, i9 CPU), which lets it answer complex neoantigen prediction queries in real time. In short, DeepPROTECTNeo has a unified, reproducible, and fully documented training pipeline that includes strict folding, standardized preprocessing, strong negative sampling, and the latest deep learning optimization. These strategies are what make the model's predictions strong, stable, and useful in a wide range of clinical settings. They also make it easy to use as both a research framework and a web service for the wider immuno-oncology community.

Note S3. Performance Evaluation Metrics

Based on the SoftMax scores provided by each model, we employed multiple evaluation metrics, each capturing different aspects of classification quality for assessing model performances. Balanced accuracy was used as a primary measure because it equally accounts for class imbalance by averaging sensitivity (true positive rate) and specificity (true negative rate), thereby providing a more reliable assessment than raw accuracy in skewed datasets:

$$\text{ Balanced Accuracy }=\frac{1}{2}\left( \frac{TP}{TP+FN}+\frac{TN}{TN+FP} \right),$$

(1)

We further report the Area Under the Receiver Operating Characteristic Curve (AUROC), which evaluates the model's ability to discriminate between classes across thresholds, and the Area Under the Precision-Recall Curve (AUPRC), particularly informative for imbalanced settings. Additionally, we compute standard classification metrics, including precision (the proportion of predicted positives that are correct), recall (the proportion of actual positives correctly identified), and their harmonic mean, the F1 score. For multi-class settings, we report the macro-averaged F1 score, which equally weights each class regardless of frequency. The formulas for these metrics are as follows:

$$\text{ Precision }=\frac{TP}{TP+FP}, \text{ Recall }=\frac{TP}{TP+FN}, F1=2\cdot\frac{\text{ Precision }\cdot\text{ Recall }}{\text{ Precision }+\text{ Recall }},$$

(2)

The macro F1 score is computed as the unweighted average of per-class F1 scores. Together, these metrics provide a robust and balanced evaluation of model performance, capturing threshold-independent ranking ability (AUROC, AUPRC) and class-specific prediction quality (balanced accuracy, precision, recall, F1).

Note S4. TCR-Epitope Binding Score Aggregation

To estimate the immunogenic potential of each candidate epitope in the context of the patient's T-cell repertoire, we computed an epitope-level summary score based on pairwise binding predictions from our DeepPROTECTNeo model. This score, termed the TCR Specificity Score, reflects the strength of interaction between an epitope and the most reactive clonotypes within the patient's TCR repertoire.

Let $\mathcal{E}=\left\{ e_{1},e_{2},\ldots,e_{m} \right\}$ be the set of candidate peptide-MHC epitopes, and let $\mathcal{T}_{p}=\left\{ t_{1},t_{2},\ldots,t_{n} \right\}$ denote the set of TCRs derived from RNA-seq of patient $p$. For each TCR-epitope pair $\left( t_{i},e_{j} \right)$, DeepPROTECTNeo predicts a binding score $S_{ij}\in[0,1]$, where higher values indicate stronger likelihood of recognition.

For each epitope $e_{j}$, we compute the 95th percentile of its predicted binding scores across all patientspecific TCRs:

$$\text{ TCR Specificity Score }_{e_{j}}=\text{ Percentile }_{g_{5}}\left( \left\{ S_{ij}\mid t_{i}\in\mathcal{T}_{p} \right\} \right)$$

(3)

This percentile-based aggregation ensures robustness to noise and outliers while capturing strong interaction signatures representative of potentially reactive T-cell subpopulations. Unlike the maximum score, which is sensitive to noise, the 95th percentile captures a small subset of high-confidence interactions.

Note S5. Retention Threshold Calculation

The retention threshold $\tau$ was computed as the 95th percentile of TCR Specificity Scores among nonvalidated epitopes in the TESLA benchmark cohort:

$$\tau=\text{ Quantile }_{0.95}\left( \left\{ \text{ TCR Specificity }\text{ Score }_{e_{j}}\mid e_{j}\notin\mathcal{V} \right\} \right)$$

(4)

This cutoff ensures that only 5% of non-validated epitopes (i.e., potential false positives) are retained by the model, resulting in high specificity. In our dataset, this yielded a retention threshold of 0.48 .

An epitope $e_{j}$ was considered retained if:

$$\text{ TCR Specificity Score }_{e_{j}}\geq\tau=0.48$$

(5)

This strategy balances immunogenic epitope sensitivity with false positive suppression, consistent with stringent biomarker discovery practices.

Note S6. Hierarchical Clustering of Cross-Dataset Feature Importance

All descriptors were first aggregated into a matrix $R\in\mathbb{R}^{F\times3}$, where $F$ is the total number of unique features and each column corresponds to one of the three datasets; any missing entry was imputed as

$$R_{f,d}^{'}=\left\{ \begin{matrix} R_{f,d}, & \text{ if present }, \\ \max_{f^{'},d^{d}} \left( R_{f^{'},d^{d}} \right)+1, & \text{ if absent }, \end{matrix} \right.$$

(6)

to ensure every feature appears in every column. We then rescaled ranks to a $[0,1]$ importance score via

$$H_{f,d}=\frac{\left( \max R^{'}+1 \right)-R_{f,d}^{'}}{\left( \max R^{'}+1 \right)-1},$$

(7)

so that higher values indicate stronger feature importance. Next, we quantified pairwise dissimilarity between features by computing the Euclidean distance

$$D_{f,g}=\sqrt{\sum_{d=1}^{3} \left( H_{f,d}-H_{g,d} \right)^{2}}$$

(8)

and applied agglomerative hierarchical clustering using the average-linkage criterion, in which the distance between two clusters $C_{i}$ and $C_{j}$ is defined as

$$d\left( C_{i},C_{j} \right)=\frac{1}{\left| C_{i} \right|\left| C_{j} \right|}\sum_{f\in C_{i}} \sum_{g\in C_{j}} D_{f,g}.$$

(9)

The resulting dendrogram orders the feature rows so that those with similar importance profiles lie adjacent (columns remain in their fixed order). When plotted as a heatmap with this clustered row order, tight subtrees visually highlight groups of descriptors sharing coherent cross-dataset importance patterns, and branch lengths directly encode the magnitude of their dissimilarity.
