## Additional File 2 for "DeepPROTECTNeo: A Context-aware Personalized and Reverse Vaccinology-guided Deep Learning Framework for Immunogenicity Prediction"

**(Additional File 2: Supplementary Figure S1-S17)**

Pralay Mitra, Ph.D.

Department of Computer Science and Engineering,

Indian Institute of Technology Kharagpur

West Bengal - 721302, India

ORCID ID: 0000-0003-4119-3788

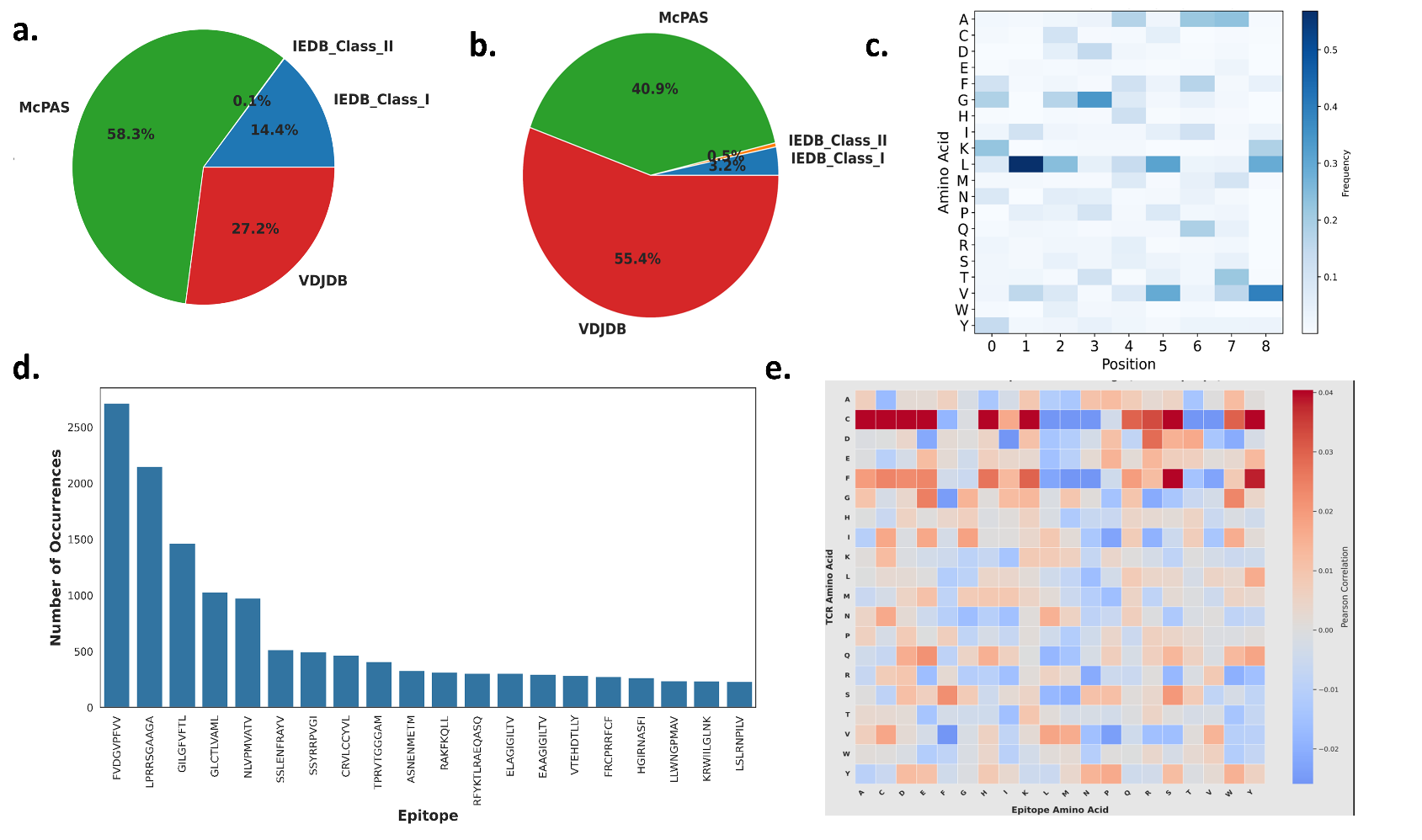

**Figure S1. Overview of the datasets used and their distribution a)** Number of unique CDR3β across all the datasets **b)** Number of unique epitopes across all the datasets c) Distribution of the Most Frequent Epitopes in the Training Dataset d) The 20 most occurring epitopes across all the datasets e) Amino acid usage correlation between TCR CDR3β and epitope sequences in the unified dataset

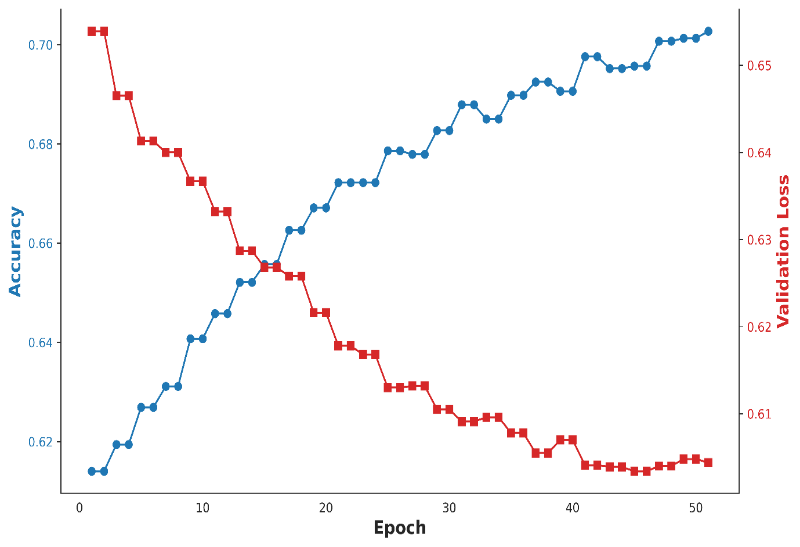

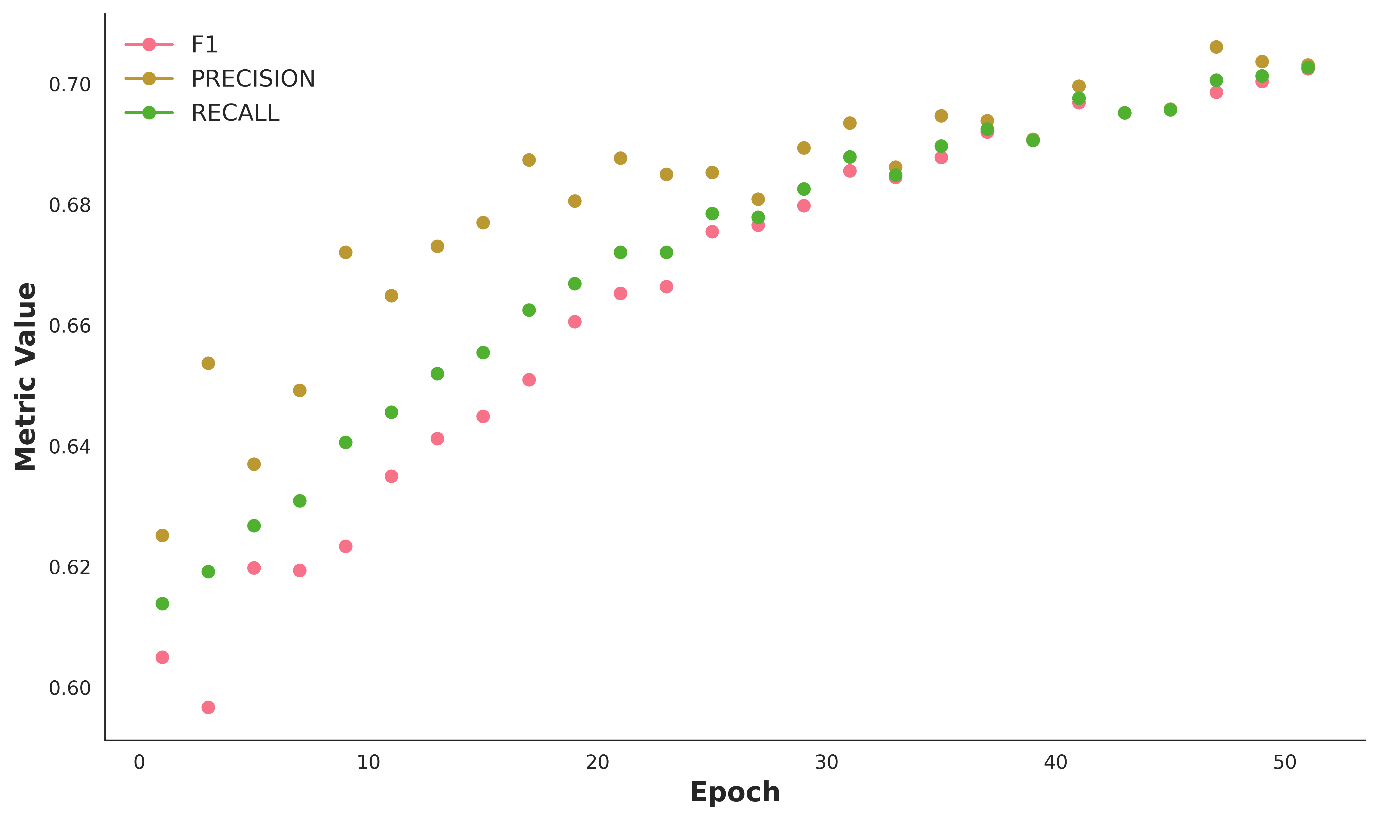

**a.**

**b.**

**Figure S2. Overview of the performance across the epochs a)** Epoch-wise evolution of training accuracy and validation loss for the optimal model hyperparameter configuration during grid search, measured on an independent 80%/20% training/validation partition. The curves demonstrate steady convergence, with rapid early gains in accuracy and a corresponding reduction in validation loss, plateauing as the model approaches optimal generalization. **b)** Longitudinal tracking of F1 score, precision, and recall over the initial 30 training epochs reveals uniform and synchronized improvement of all classification metrics, with minimal divergence between them, signifying stable model calibration and balanced performance across both positive and negative classes. These results collectively underscore the efficiency and robustness of DeepPROTECTNeo’s optimization procedure and its capacity to achieve high, stable predictive accuracy through end-to-end learning on large, complex immunological datasets.

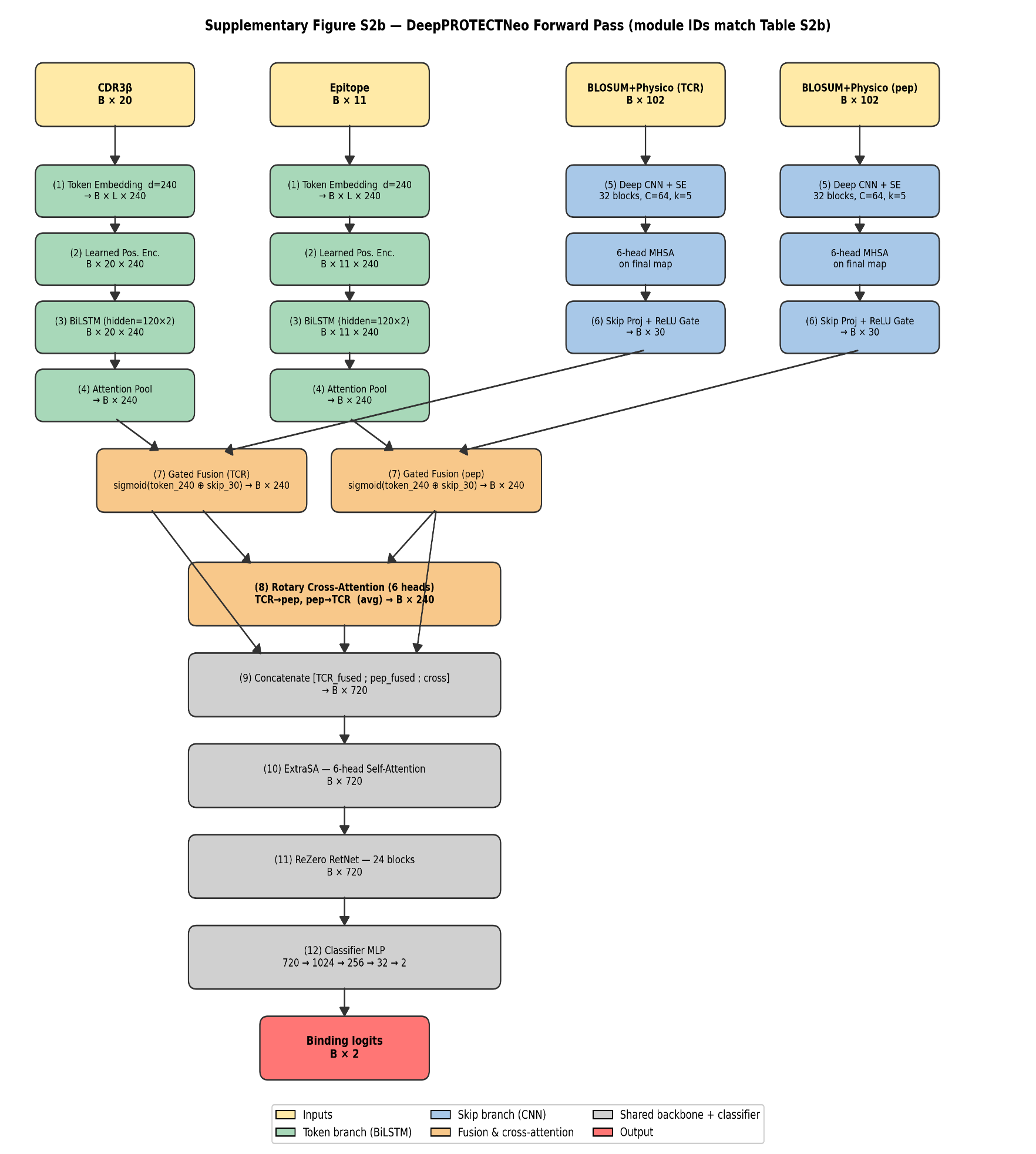

**Figure S3. Shape-annotated forward-pass block diagram of DeepPROTECTNeo.** Module IDs (1–12) correspond to rows in Supplementary Table S2. B = batch size; L = sequence length; d = d_model = 240. Dual-branch: Token Branch (modules 1–4) + Skip Branch (5–6) → Gated Fusion (7) → Cross-Attention (8) → RetNet (9–10) + ExtraSA (11) → MLP (12).

**Figure S4. The effect of the feature projector components on our architecture. a)** The scatter plot shows the relationship between the number of convolutional blocks (cnn_blocks) and the best validation loss achieved across all grid search runs **b)** The scatter plot shows the relationship between the number of convolutional blocks (skip_dim) and the best validation loss achieved across all grid search runs. **c)** The bar plot quantifies the statistical significance of each hyperparameter in explaining variation in validation loss, as measured by the ANOVA F-score. Higher F-scores indicate a greater impact on performance. The plot reveals that cnn_dim and cnn_blocks are the most influential hyperparameters, followed by skip_dim. Other parameters, such as n_heads, cnn_ch, and learning rate (lr), exhibit minimal direct effect within the tested ranges. This analysis guides prioritization for future hyperparameter optimization.

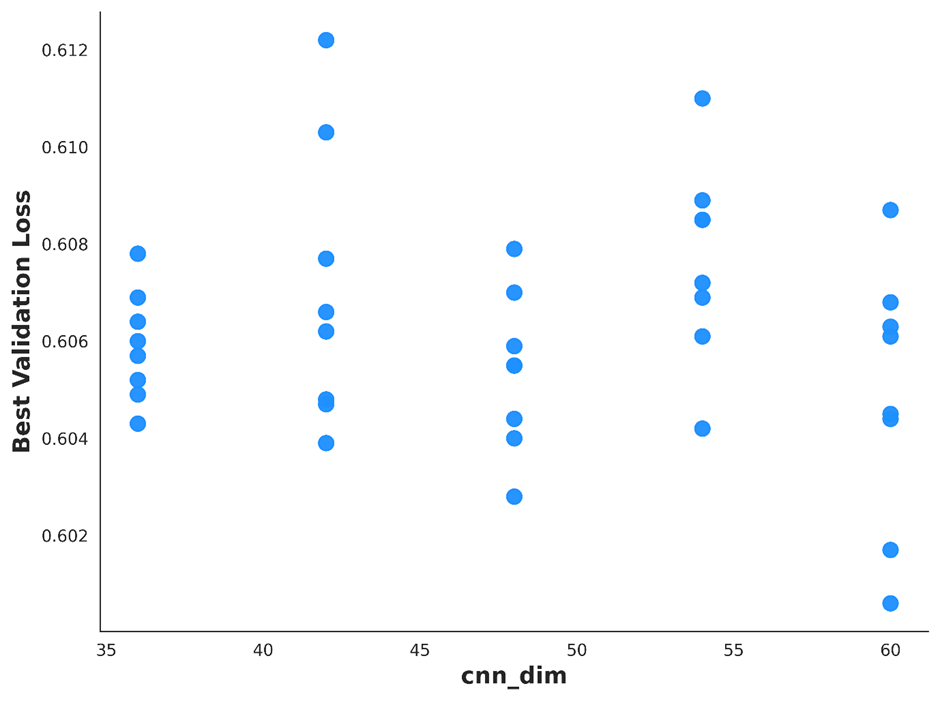

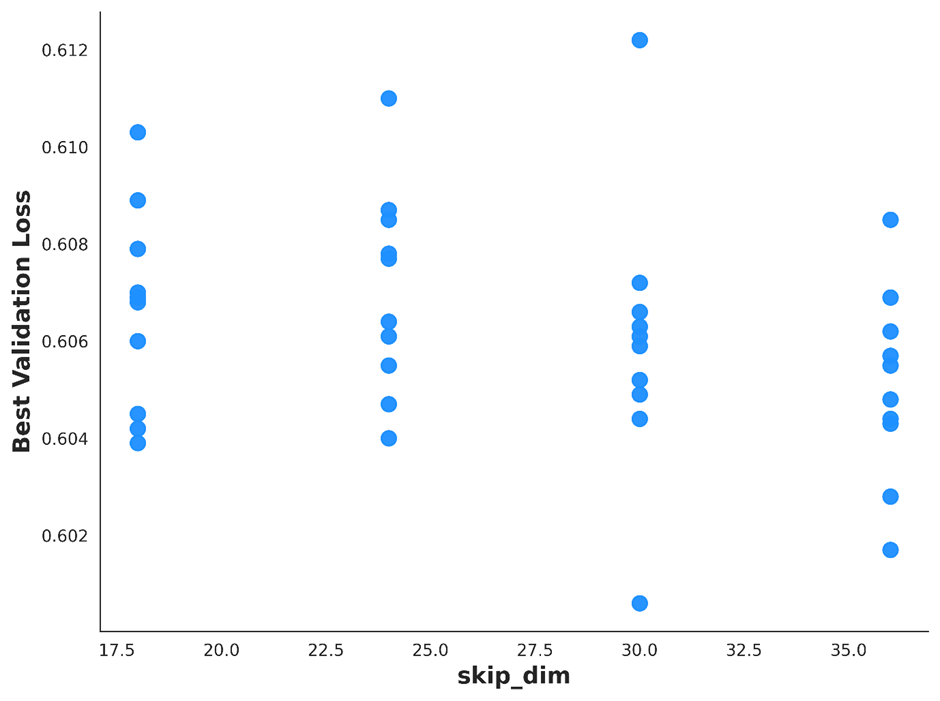

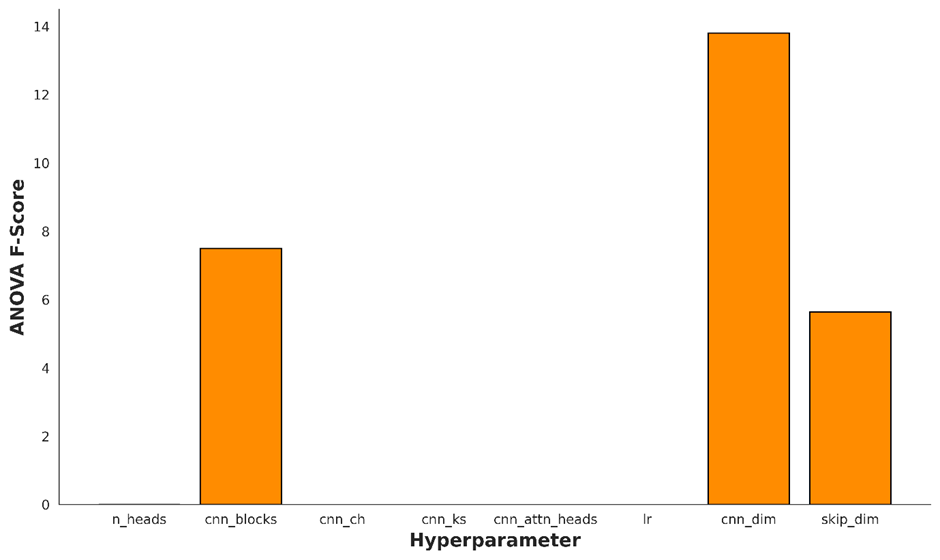

**a.**

**b.**

**c.**

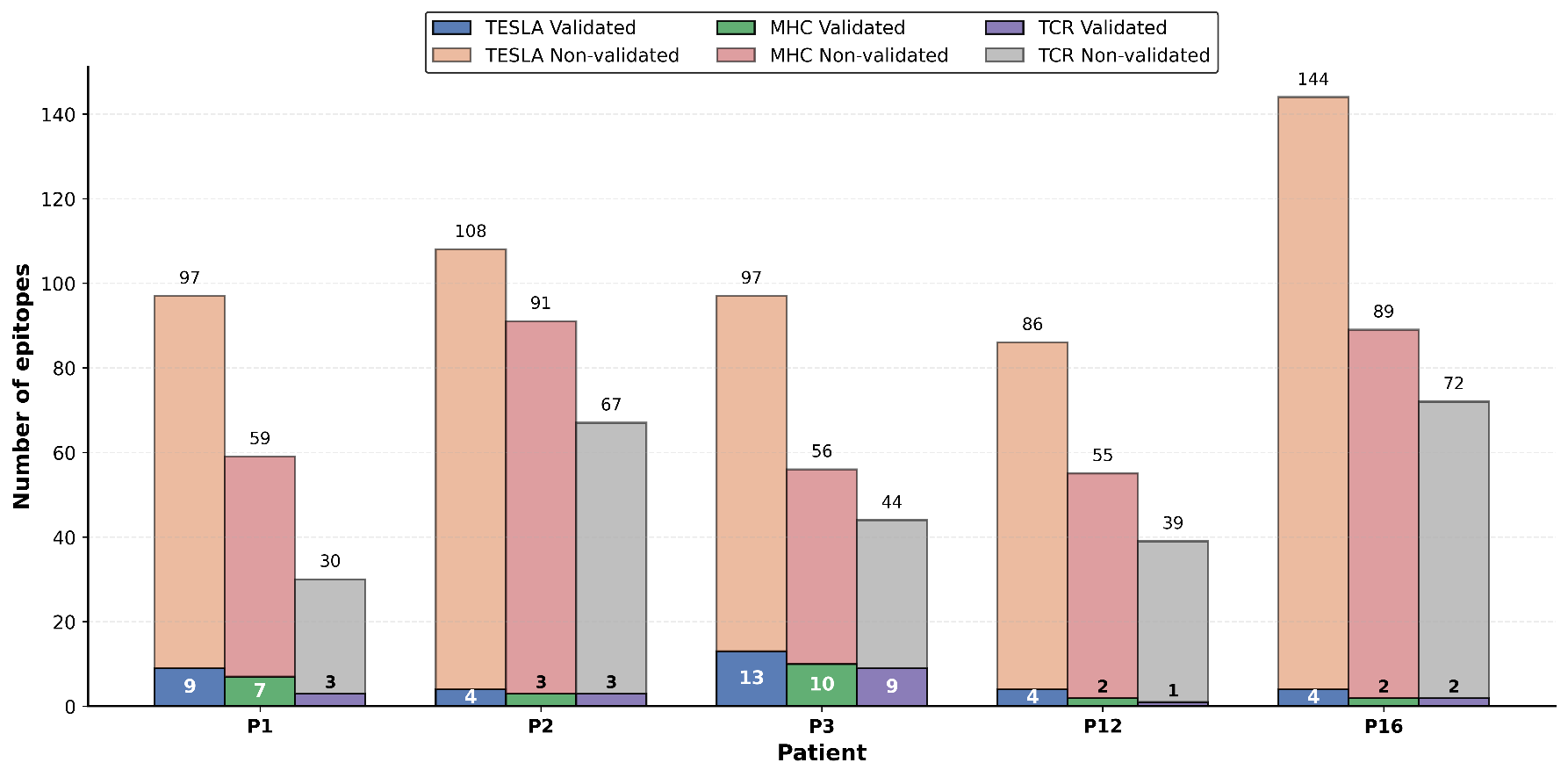

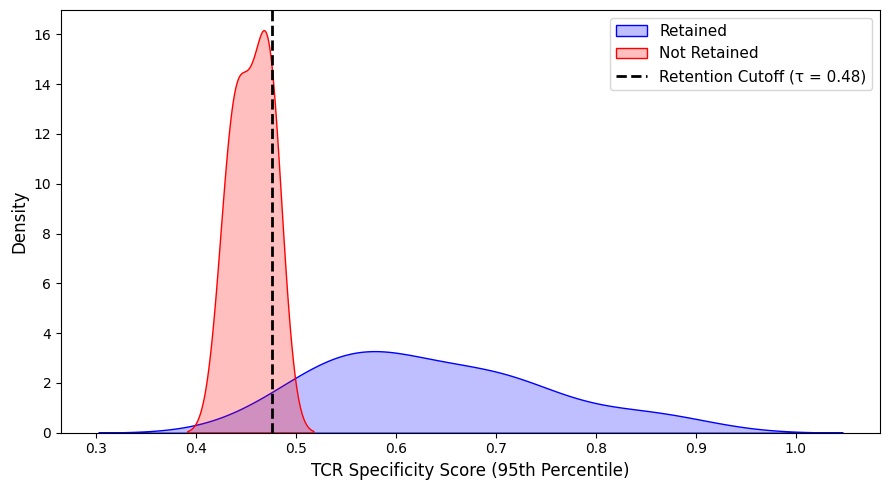

**Figure S6**. **Illustrating the distribution of TCR specificity scores (95th percentile) across retained and non-retained epitopes.** The retention threshold τ = 0.48, derived as the 95th percentile of scores from the non-validated epitope pool, effectively separates validated (retained) epitopes from the background, as shown by the non-overlapping density distributions.

**Figure S5. Per-patient comparison of epitope counts from the TESLA consortium versus our MHC- and TCR-binding predictions**. For each patient (IDs 1, 2, 3, 12, 16), three adjacent bars show: TESLA-derived epitopes (blue/orange), predicted MHC binders (green/red), and predicted TCR binders (purple/gray). Within each bar, the solid (darker) portion denotes **validated** epitopes and the translucent (lighter) portion denotes **non-validated** epitopes. The number printed immediately above each solid segment is the count of validated epitopes, and the number above the full stack is the total epitope count for that category.

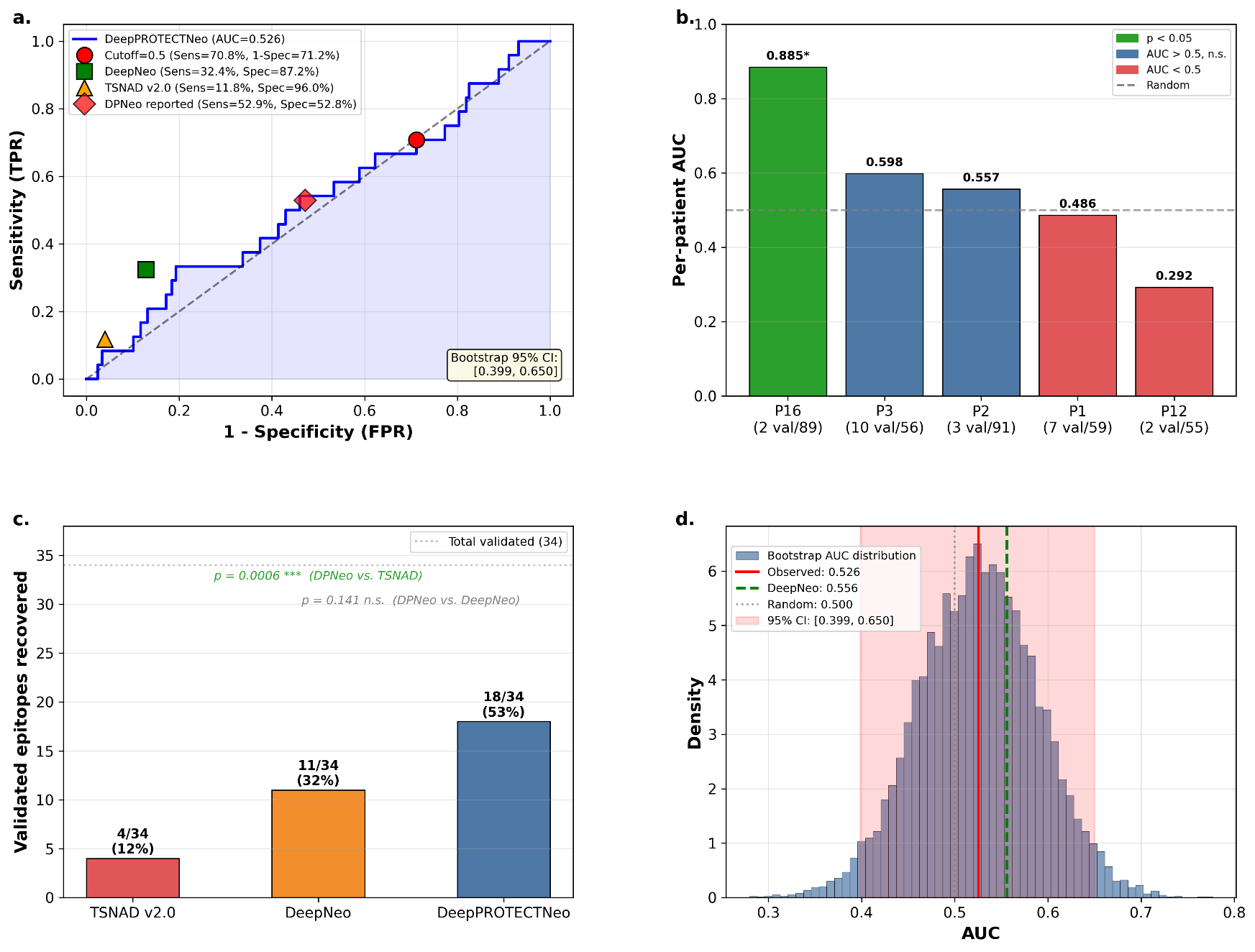

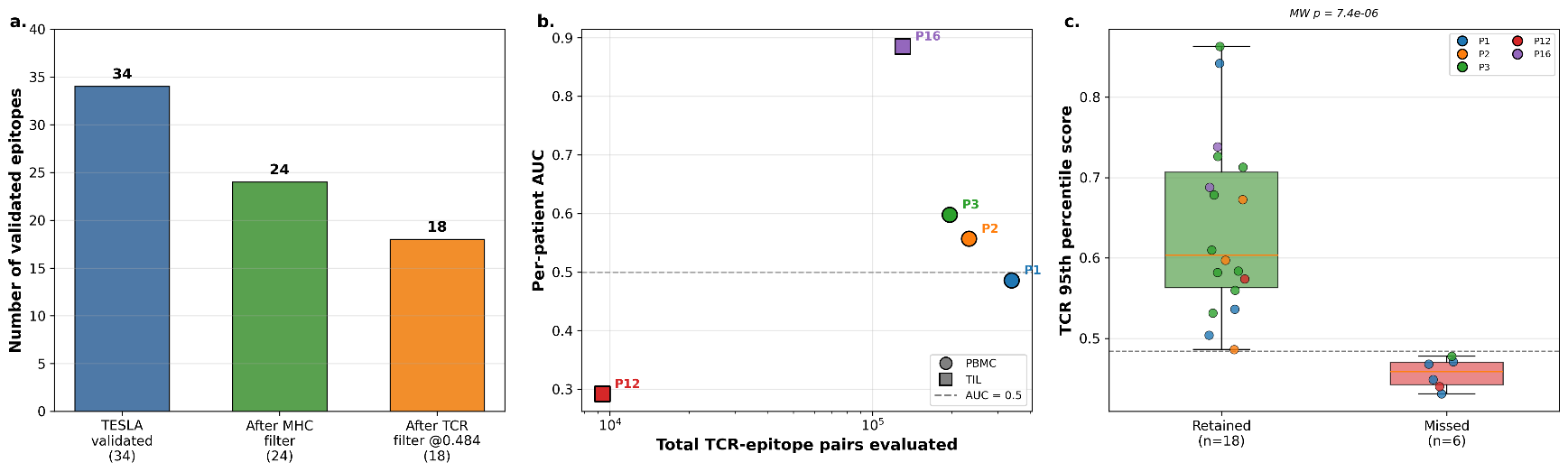

**Figure S7. Missed epitope analysis for the TESLA cohort.** **(a)** Stage-wise survival of the 34 TESLA-validated immunogenic epitopes through the DeepPROTECTNeo pipeline: 24 retained after the MHC filter, 18 after the TCR filter at the data-derived cutoff of 0.484. **(b)** Per-patient AUC vs. total number of TCR–epitope pairs evaluated (log scale); circles = PBMC, squares = TIL. **(c)** TCR 95th-percentile score distributions for validated epitopes retained (n = 18) vs. missed (n = 6); Mann–Whitney p = 7.4 × 10⁻⁶.

**Figure S8. Per-patient AUC analysis.** **(a)** Per-patient AUC values with bootstrap 95% CI. **(b)** Score distributions (validated vs. non-validated epitopes) as violin plots. **(c)** Per-patient ROC curves with permutation p-values. Patient 16 (AUC = 0.885, p = 0.032) demonstrates strong discrimination. **(d)** Bootstrap (1000 resamples) of the pooled TESLA AUC for DeepPROTECTNeo (observed = 0.526; 95 % CI [0.399, 0.650]) compared against DeepNeo (0.556) and the random baseline (0.500).

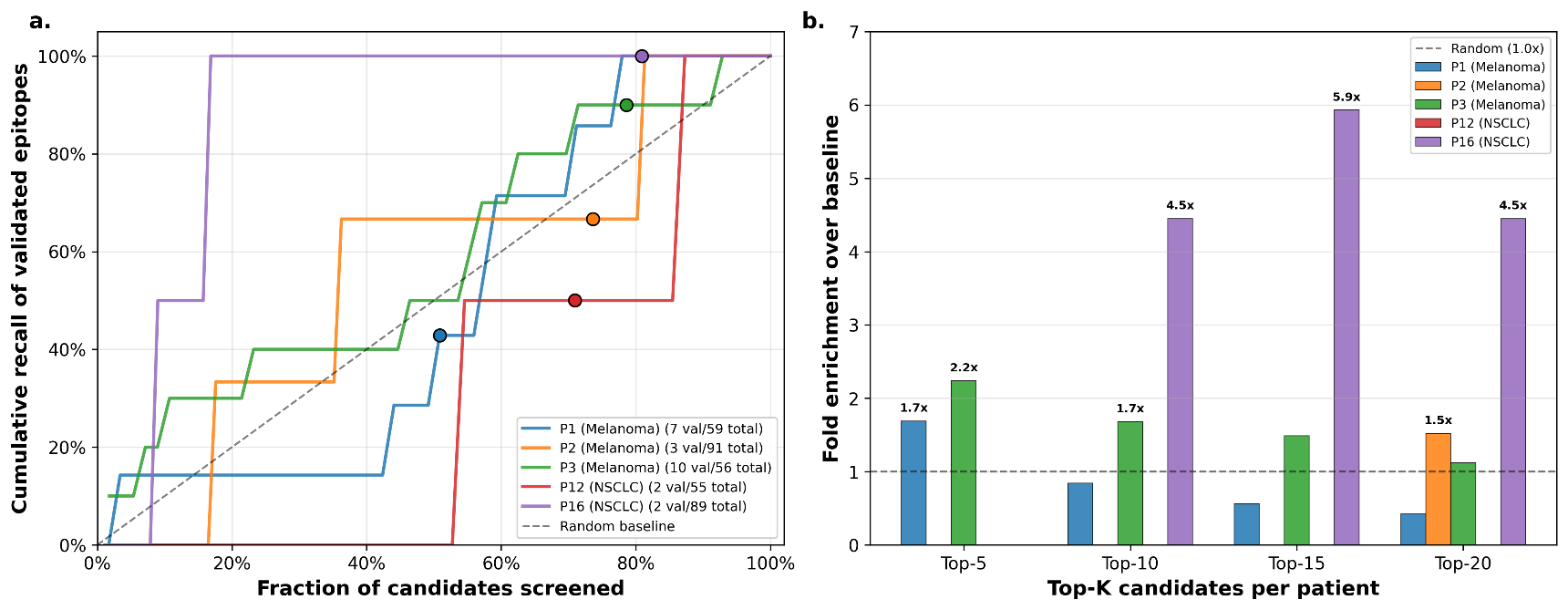

**Figure S10.** **Integrating TCR Context Substantially Improves Neoantigen Prioritization. (a)** Receiver operating characteristic (ROC) curves comparing logistic regression (blue) and random forest (green) models trained on TESLA epitope features, evaluated via 5-fold cross-validation. Logistic regression achieved superior performance (AUC = 0.81) relative to random forest (AUC = 0.72), indicating robust linear separability of validated versus non-validated epitopes. **(b)** Feature importance expressed as odds ratios from the logistic regression model. The TCR 95th percentile specificity score was the strongest predictor (odds ratio ≈ 1.5), followed by foreignness (≈1.3) and antigenic processing markers (agreptopicity, binding stability). These results underscore the critical contribution of patient-specific TCR repertoire context to neoepitope prioritization, complementing classical antigen presentation features.

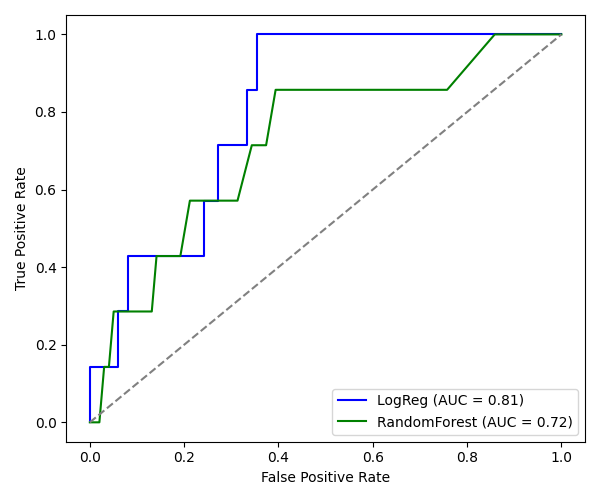

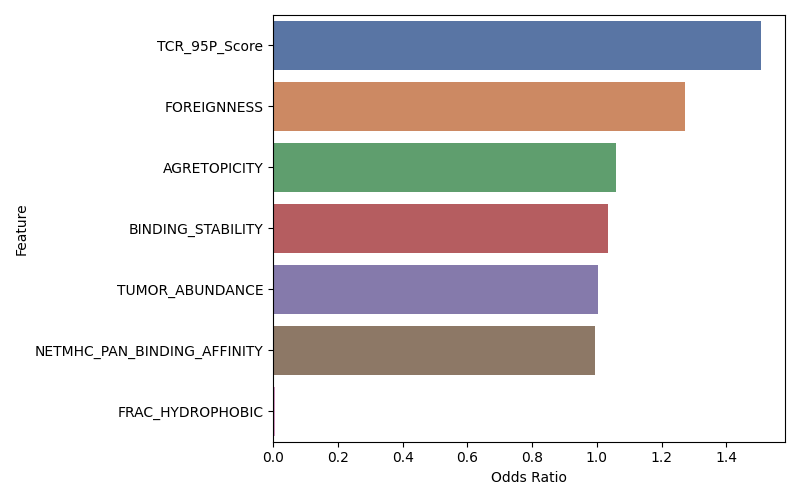

**a.**

**b.**

**Figure S9. Per-patient ranking performance on the TESLA cohort: cumulative recall and Top-K fold-enrichment of validated epitopes. (a)** Cumulative recall of validated epitopes vs. fraction of candidates screened, per patient; dashed line = random baseline; markers mark the 0.5-cutoff operating point. **(b)** Fold enrichment over the per-patient baseline immunogenicity rate at Top-K = 5, 10, 15, 20; dashed line = random (1.0×).

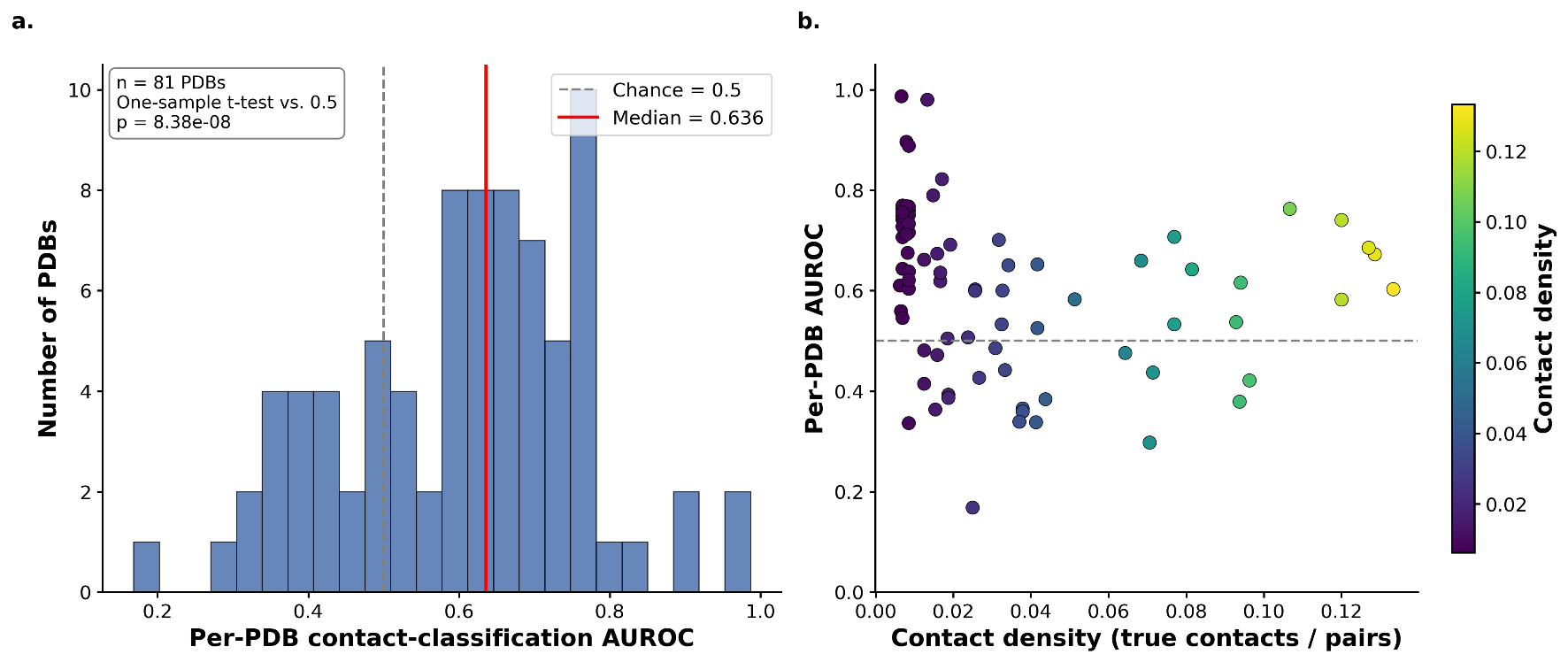

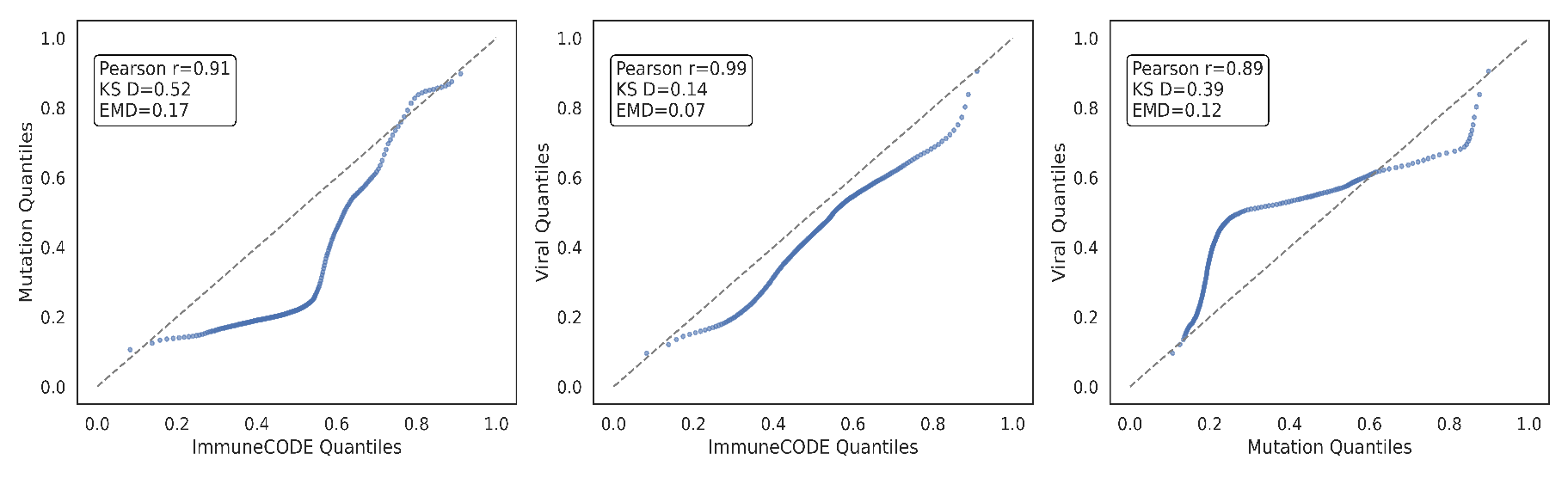

**Figure S12.** **Structural validation of attention-based contact prediction across 81 TCR–pMHC crystal structures (per-PDB AUROC histogram; AUROC vs. contact density).** **(a)** Histogram of per-PDB contact-classification AUROC across 81 crystal structures. Median = 0.636; one-sample t-test p = 8.4 × 10⁻⁸ vs. random baseline (0.5, dashed line). **(b)** Per-PDB AUROC plotted against contact density (fraction of CDR3β residues making peptide contacts)

**Figure S11. Scatter plots showing pairwise comparisons of peptide-descriptor importance ranks between (left) ImmuneCODE vs. Mutation, (middle) ImmuneCODE vs. Viral, and (right) Mutation vs. Viral**. Each point represents a single descriptor, plotted by its rank in one dataset (x-axis) against its rank in the other (y-axis); lower ranks indicate higher importance. The dashed diagonal line marks perfect agreement (y = x). Inset annotations give Spearman’s rank correlation (ρ) and two-tailed p-value for each comparison, demonstrating strong overall concordance (ρ = 0.94, 0.92, 0.90; p < 10⁻³⁸) and a small number of outliers that reveal dataset-specific feature weighting.

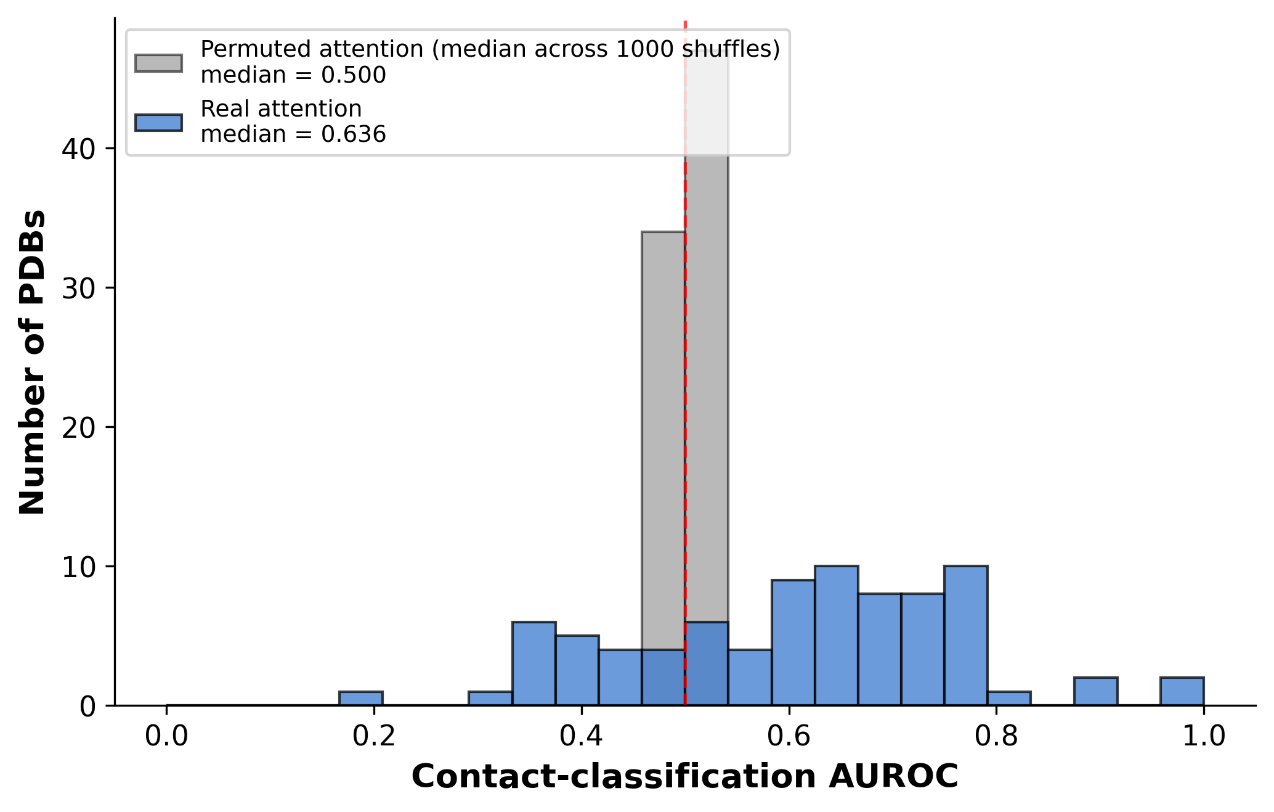

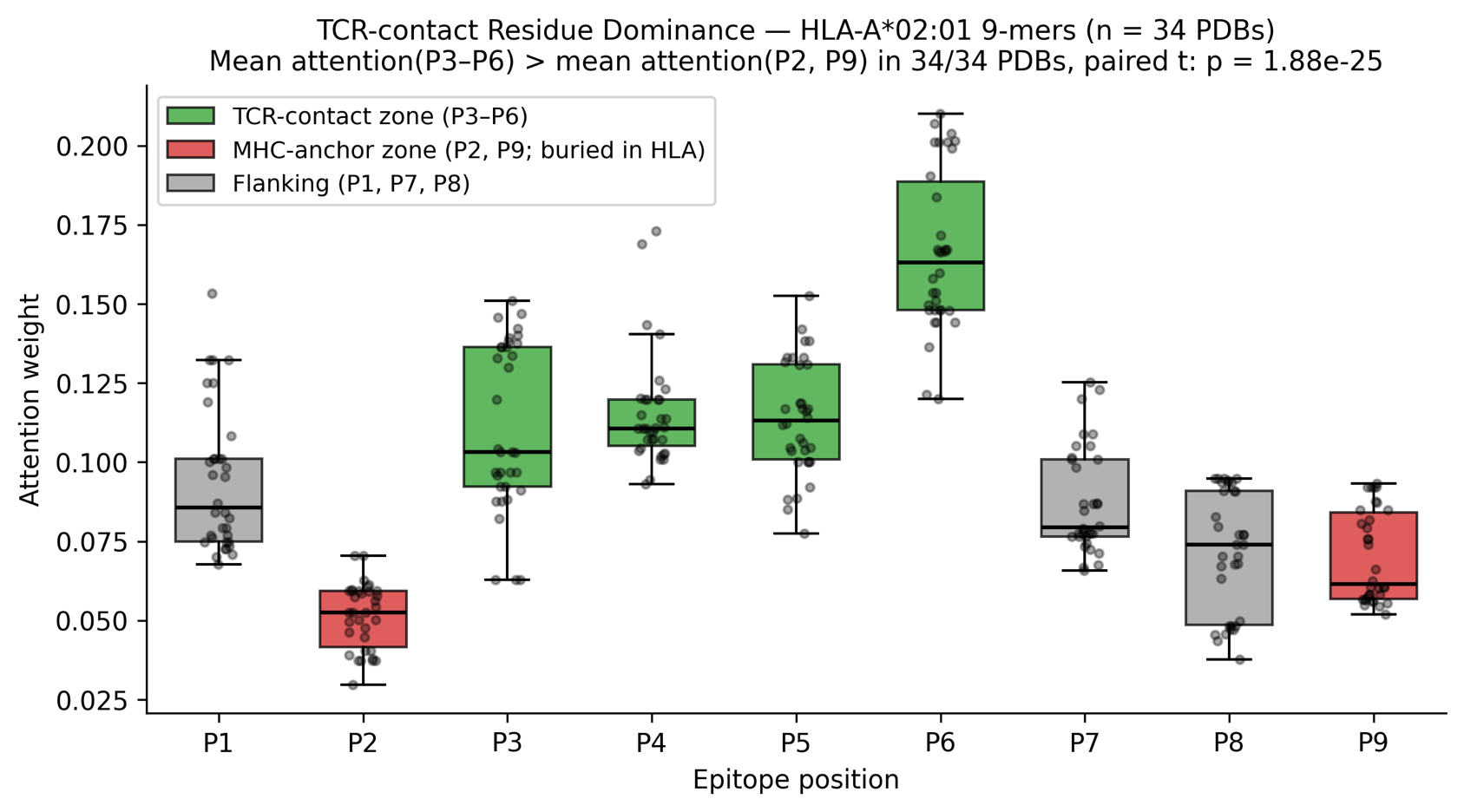

**Figure S14.** **Per-position mean attention (P1–P9) across 34 HLA-A*02:01 9-mer PDBs (±95% CI). Red = TCR-contact zone (P3–P6); grey = MHC-buried anchors (P2, P9). Mean: 0.1265 vs. 0.0606 (2.09-fold; p = 1.9 × 10⁻²⁵).**

**Figure S13.** **Real vs. permuted AUROC distributions across 81 PDBs. Real median 0.636 significantly exceeds permuted median 0.500 (paired t-test p = 1.0 × 10⁻⁷).**

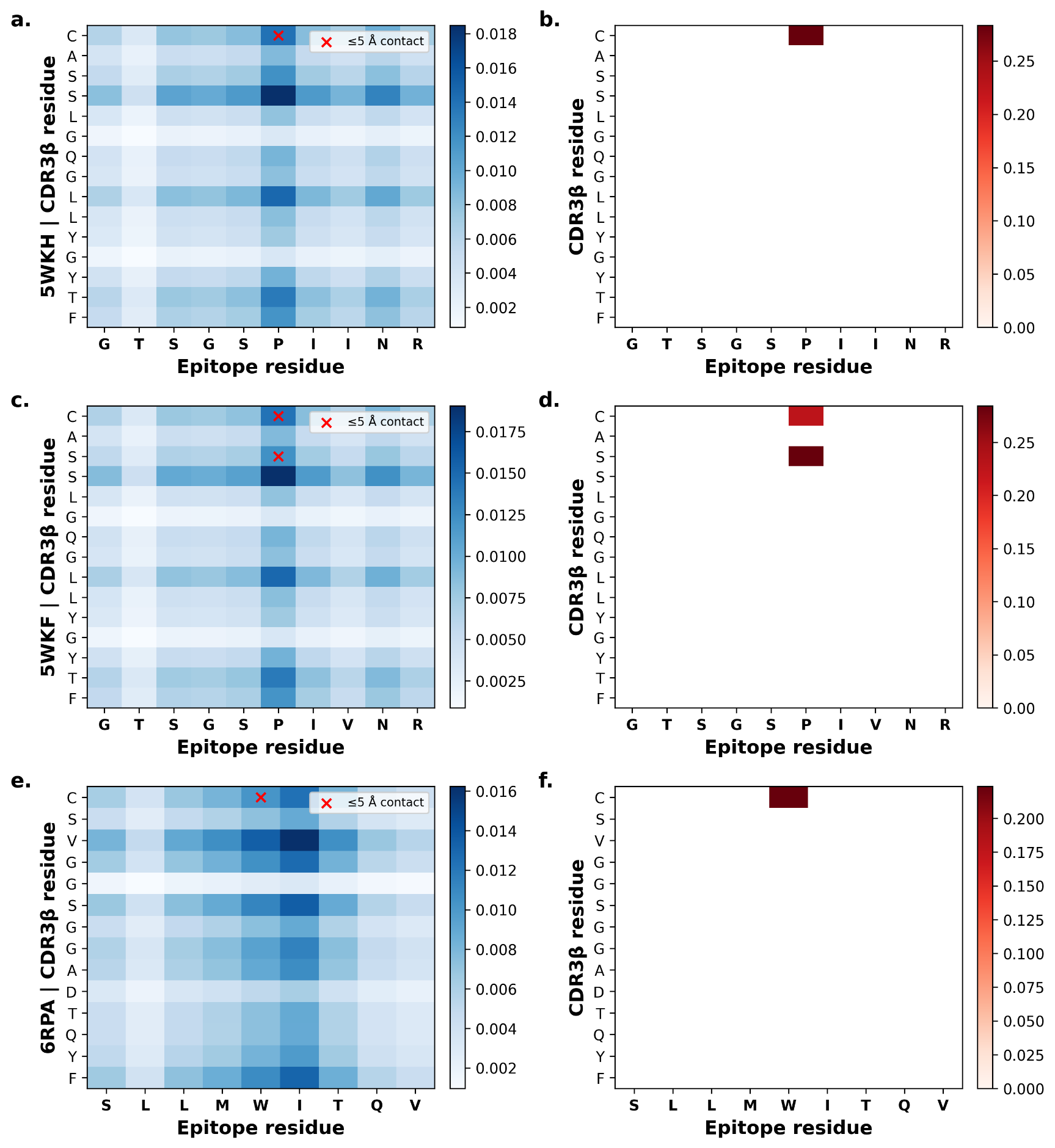

**Figure S15.** **Side-by-side attention (a) and contact (b) visualisations for the three highest-AUROC PDBs (5WKH, 5WKF, 6RPA). Left = attention heatmap A[i,j]; right = experimental contact map (≤5 Å).**

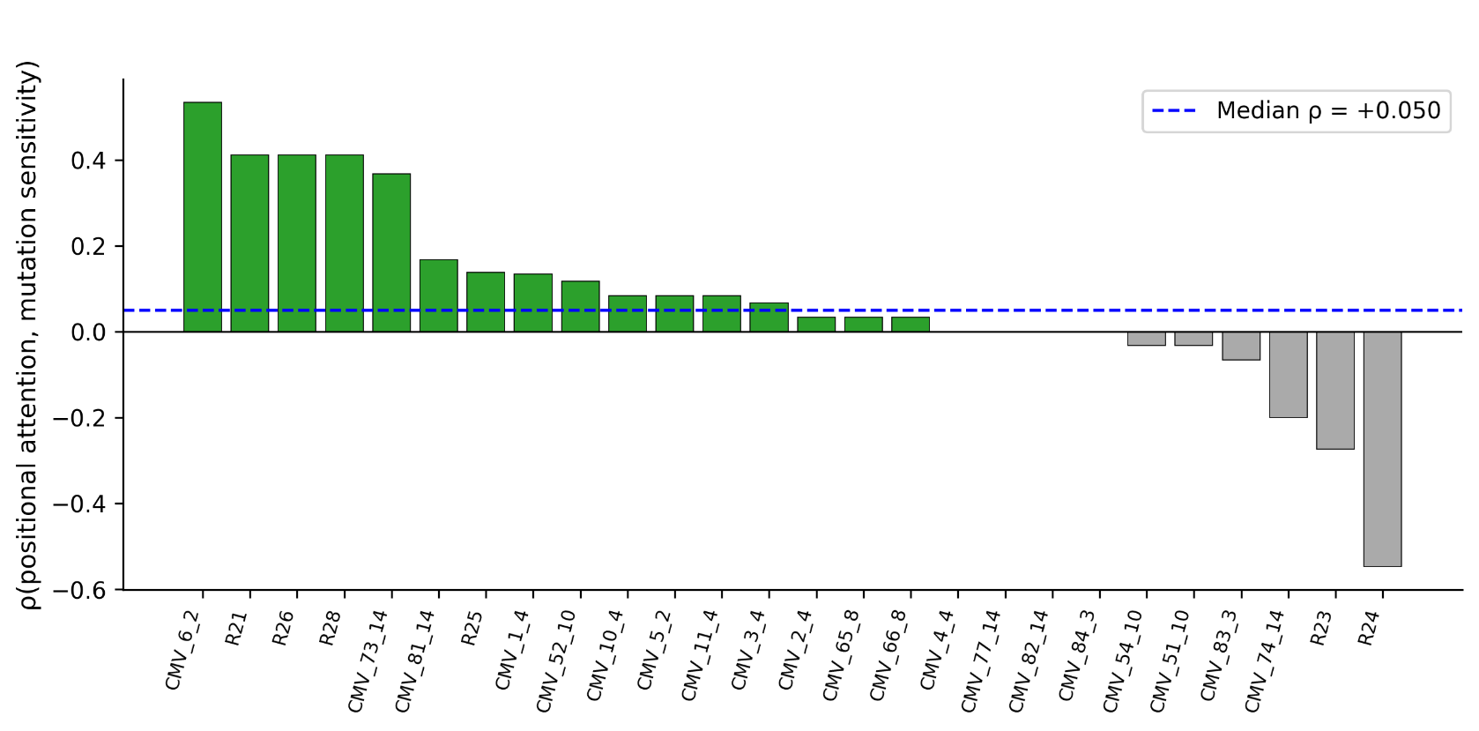

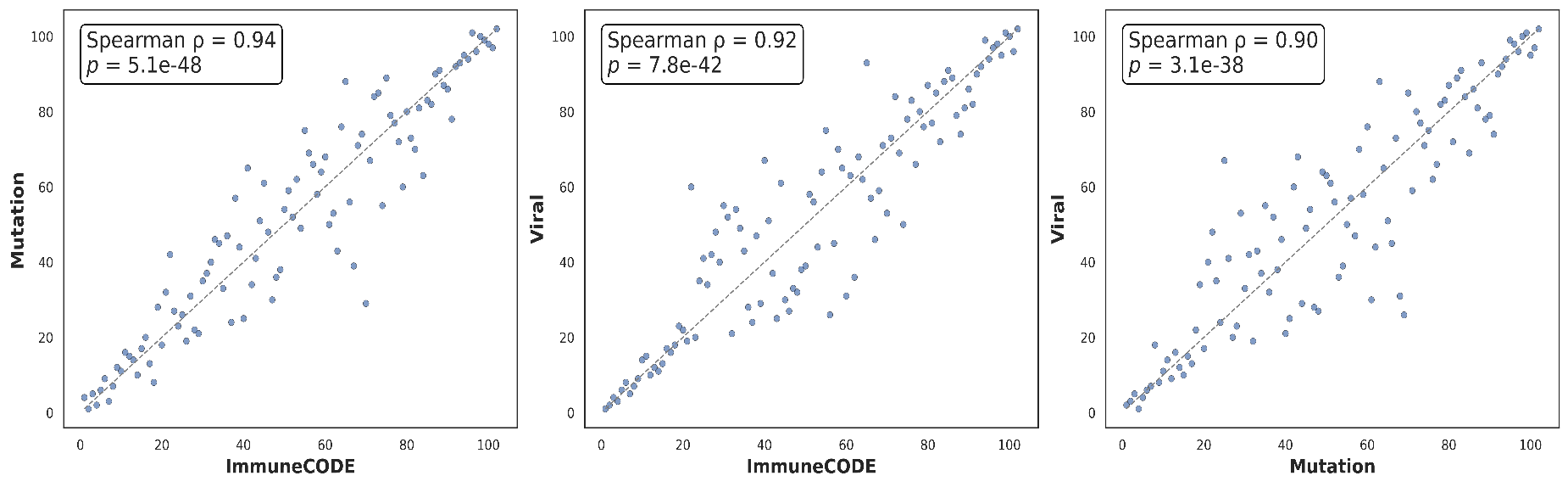

**Figure S17. Quantile–Quantile Comparison of DeepPROTECTNeo Predictions Across Independent Datasets**. Each panel displays a Q–Q plot comparing the empirical percentiles (0th–100th) of DeepPROTECTNeo’s pooled prediction scores between two independent test sets: ImmuneCODE vs Mutation (left), ImmuneCODE vs Viral (center), and Mutation vs Viral (right). Points lie on the dashed diagonal when the two distributions perfectly agree at that quantile. Inset text shows the Pearson correlation (*r*) between the quantile series (assessing linear concordance), the two‑sample Kolmogorov–Smirnov statistic D and its *p*-value (testing for any overall distributional difference), and the Earth Mover’s Distance (EMD; quantifying the average magnitude of score shifts). High *r* values indicate strong median alignment, whereas larger KS D and EMD values reveal heavier tails in Mutation and a more centrally compressed range in Viral compared to ImmuneCODE.

**Figure S16.** **Per-TCR attention vs. mutation-sensitivity scatter across 26 CDR3β clonotypes. Spearman ρ annotated per panel. Blue = ρ > 0; grey = ρ < 0. Wilcoxon p = 0.024.**
